# Instant Assembly of Collagen for Scaffolding, Tissue Engineering, and Bioprinting

**DOI:** 10.1101/2023.10.08.561456

**Authors:** Xiangyu Gong, Zhang Wen, Zixie Liang, Hugh Xiao, Sein Lee, Thomas Wright, Ryan Y. Nguyen, Alejandro Rossello, Michael Mak

## Abstract

Controllable assembly of cells and tissues offers potential for advancing disease and development modeling and regenerative medicine. The body’s natural scaffolding material is the extracellular matrix, composed largely of collagen I. However, challenges in precisely controlling collagen assembly limit collagen’s applicability as a primary bioink or glue for biofabrication. Here, we introduce a set of biopatterning methods, termed Tunable Rapid Assembly of Collagenous Elements (TRACE), that enables instant gelation and rapid patterning of collagen I solutions with wide range of concentrations. Our methods are based on accelerating the gelation of collagen solutions to instantaneous speeds via macromolecular crowding, allowing versatile patterning of both cell-free and cell-laden collagen-based bioinks. We demonstrate notable applications, including macroscopic organoid engineering, rapid free-form 3D bioprinting, contractile cardiac ventricle model, and patterning of high-resolution (below 5 (m) collagen filament. Our findings enable more controllable and versatile applications for multi-scale collagen-based biofabrication.

## Introduction

Collagen I is a prominent extracellular matrix (ECM) protein that scaffolds cells in tissues. It provides structural support and biophysical cues to govern cell behavior. Collagen-based scaffolds are widely used in tissue engineering, owing to their natural biocompatibility and to their promotion of bioactivity^1–4^. Various methods with different gelling conditions have been developed to tune collagen network properties, including pore size, fiber geometry (e.g. thickness, length), topography, and mechanics^5–12^. However, standard methods of collagen fabrication have limited versatility and tunability, as the self-assembly of collagen solutions into fiber networks is difficult to precisely control spatially and temporally. In particular, neutralized collagen solutions (at typical working concentrations) usually require tens of minutes to fully undergo gelation^13,14^, a barrier to rapid patterning and bioprinting applications, especially applications requiring accurate and stable positioning of cells and ECM.

Recent advances in bioprinting collagen I include the usage of a granular hydrogel support bath to help maintain collagen positioning after extrusion^15–17^. The neutral-pH support bath promotes gelation of high concentration acid solubilized collagen upon extrusion, enabling high-resolution printing of complex and mainly cell-free 3D constructs^15^. The acidity promotes collagen solution fluidity before extrusion, and the high collagen concentration enables fast gelation upon neutralization in the support bath, which together facilitate collagen printability. However, high-concentration collagen restricts cell infiltration, migration, and self-organization^18,19^ due to exceedingly small pore sizes, and incorporating cells in acidic collagen solutions before printing induces cell stress. Rapid patterning of cell-laden collagen inks with highly tunable features (including concentration, especially low concentration, and microarchitecture, which modulate cell signaling^5^) remains challenging. Controllable production of active and living tissues enables many applications in tissue and organoid engineering and regenerative medicine^20^.

Here, we present a new set of versatile patterning methods for unmodified collagen type I, based on accelerated collagen gelation kinetics via macromolecular crowding (MMC). Our methods, termed TRACE, enable a variety of collagen scaffolds and patterns, both cell-free and cell-laden, to be rapidly synthesized. Moreover, our methods enhance free-form bioprinting capabilities and provide enabling features, including the usage of acidic, neutralized, or cell-laden collagen solutions as bioinks; rapid bioink solidification for a wide range of collagen concentrations, unrestricted to only high concentrations (tested from 2mg/mL to 9.5mg/mL); tunable local and mesoscopic collagen architectures; and improved bioprinting resolution of collagen down to the single-digit micrometer scale.

### Instant collagen gelation into basic patterns

Macromolecular crowding (MMC) via the addition of high molecular weight PEG (polyethylene glycol 8,000 (Da)) enables instantaneous gelation of collagen (Fig. 1a). As demonstrated by shear rheology during the gelation process of neutralized collagen at room temperature, introducing a PEG solution prior to collagen solidification leads to a sudden ramping in the shear modulus, which indicates rapid gelation (Fig. 1b). Additionally, when dispensing microliter neutralized collagen droplets onto a PEG8000 bath (Fig. 1c, Extended Data Fig. 1a, Movie 1), we find the droplets rapidly and consistently gel into millimeter-scale disks with a defined boundary within 1 second (Fig.1d,e and Extended Data Fig. 1b). The boundary of the instantly formed disk slightly expands, and the disk becomes increasingly opaque from the edge to the center, reaching a plateaued opacity in approximately 3 minutes. In contrast, collagen droplets dispensed onto a PBS bath instantly disappear and only start gelling after 5 minutes into undefined shapes (Movie 2). Furthermore, the area of the microliter collagen disk can be controlled linearly by adjusting the droplet volume (Fig. 1f). High-resolution confocal imaging reveals that the disks exhibit a smooth surface, consisting of a thin, densely assembled collagen layer that rapidly forms upon contacting the PEG bath. Collagen disks containing mouse mesenchymal stem cells (mMSCs) exhibit gel compaction, spontaneous tissue folding, and osteogenic phenotype upon osteogenic induction (Extended Data Fig. 2). This provides a high-throughput application for studying cell mechanics, tissue morphogenesis, and stem cell differentiation.

**Fig. 1.**
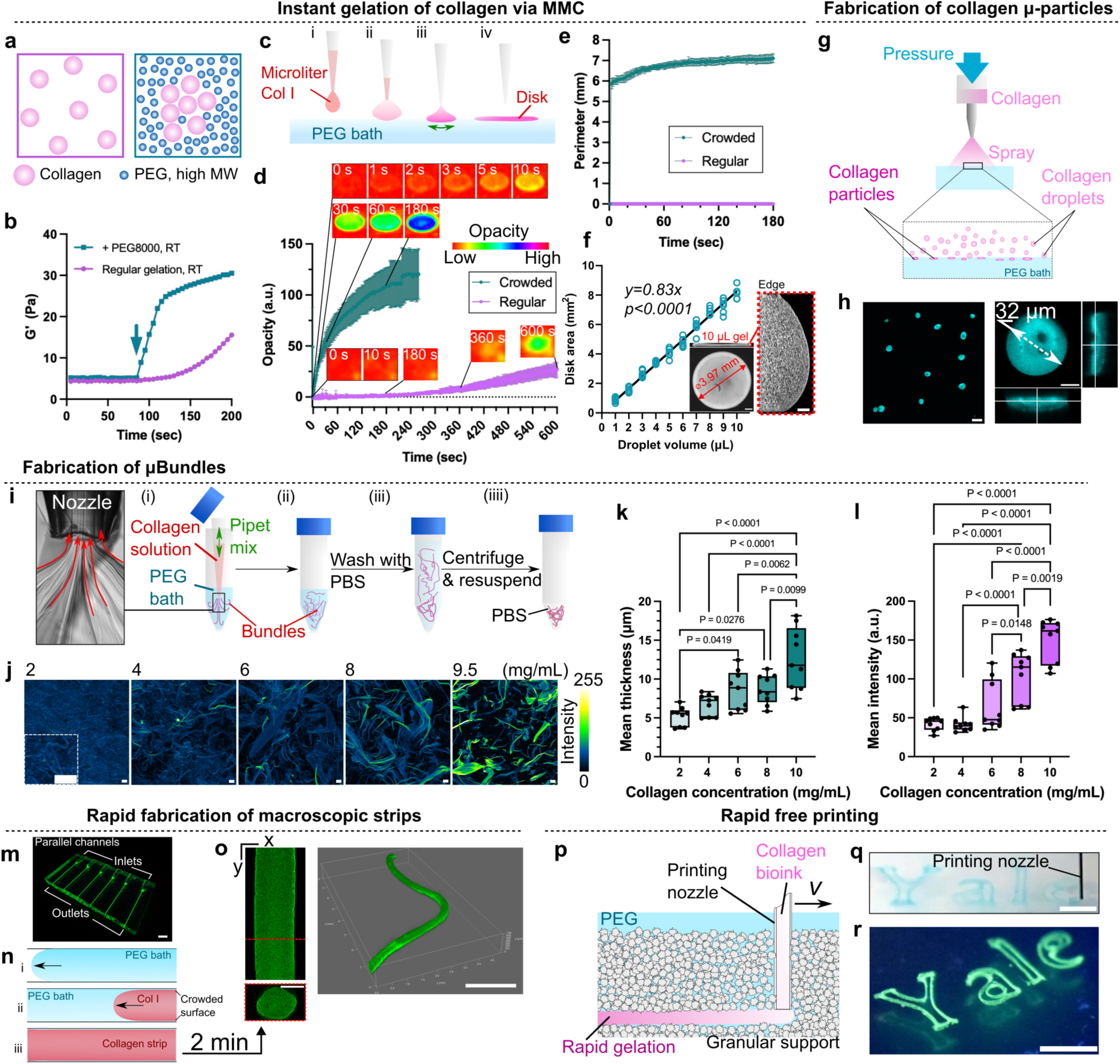
MMC-induced instant gelation facilitates rapid, multiscale collagen fabrication. (**a**) Schematic depicting the rapid assembly of collagen by the MMC agent PEG 8000. (**b**) Storage modulus (G’) of neutralized collagen upon the addition of PEG 8000 vs. the gelation at room temperature. (**c**) Schematic of collage disk formation from μ-liter collagen droplets on the surface of the PEG bath. (**d**) Collagen droplets boundary formation and gelation speed in the PEG bath compared to those in the PBS bath, as indicated by gel opacity analysis (**e**) Perimeter measurements of the rapidly emerging disks as a function of time for 3 minutes, compared with collagen droplets dispensed in PBS. (**f**) μ-liter collagen disk area as a function of the volume of collagen droplets. The relation is fitted into a linear trend. The inset image is a confocal image showing the top and side view of a 10-μL collagen disk. The close-up view enclosed in a red frame shows a distinct dense boundary formed at the edge of the disk as a result of the rapid polymerization. Scale bars, 20μm. (**g**) Schematic of collagen μ-particle fabrication via pressurized spraying method. (**h**) Fluorescent images showing the 3D morphology of collagen μ-particles fabricated with acidic 10mg/mL collagen solution. Scale bars, 50μm (left); 10μm (right). (**i**) Four-step procedure for collagen μ-bundle fabrication, using spontaneous gelation of collagen jet streams induced by fast pipet mixing within the PEG bath. The streamlines created by mixing the two liquids are indicated by red arrows in the bright field microscopic image. (**j**) Fluorescent intensity heat map indicating the final density of collagen μ-bundles fabricated at different collagen precursor concentrations (2-9.5mg/mL). Scale bars, 30μm. The images were used to characterize (**k**) bundle thickness and (**i**) bundle density as a function of varying starting collagen solution concentrations (n=9 field of view from N=3 independent samples). (**m**) Parallel PDMS channels (diameter: 400 μm) for rapid, high throughput strip production. (**n**) Schematic of a single collagen stripe production within the PEG solution-filled PDMS channel. (**o**) Fluorescently labeled collagen strip (2mg/mL) extracted from the PDMS channel 2min after fabrication. Scale bar in the cross-sectional view, 200 μm; Scale bar in the 3D reconstruction, 2mm. (**p**) Schematic of freeform bioprinting in a PEG-contained granular support bath, termed “TRACE printing bath.” (**q**) Printing multilayered 2D "Yale” logo in TRACE printing bath with neutralized 2mg/mL collagen bioink mixed with blue food color and (**r**) fluorescent microbeads. Scale bars in (**q**) and (**r**), 10mm.

The density of the collagen disks can be tuned by the concentration of collagen solution used to form the droplets (Extended Data Fig. 1d). Additionally, the MMC-induced collagen disks demonstrate an approximately two-fold increased concentration compared to regular collagen gels (gelled at 37℃ for 40min without PEG) (Extended Data Fig. 1e). Notably, we find that even acidified collagen droplets can be rapidly gelled into disks by the PEG bath, whereas no gel formed in regular gelation conditions (Extended Data Fig. 1d). Collagen gel droplets can be further downscaled and efficiently generated via air spraying (Fig. 1g). Sprayed aerosols of acidic collagen solution instantly form micro-gels upon contact with PEG solution, with micro-gel size easily reaching the scale of cells (Fig. 1h). Using this MMC-facilitated instant collagen gelation method, we next demonstrate several foundational and generalizable techniques for rapid patterning of unmodified collagen I solutions into diverse and versatile forms.

MMC facilitates the rapid generation of mesoscopic collagen fiber bundle networks. By pipet-mixing collagen solution into a PEG bath (Fig.1i), precipitated collagen streams are observed. The collagen streams are spontaneously nucleated into mesoscale collagen bundles by PEG, which can then be collected through centrifugation for tissue culture applications. We term these thick fiber bundles as “μ-bundles”. We demonstrate the tunability of the μ-bundle thickness and density by adjusting the collagen concentration of the pre-bundle collagen solution (Fig.1j-l). Higher collagen concentrations result in thicker and denser μ-bundles. This method enables the generation of substantially larger collagen bundles compared to the sub-micron collagen fibers in regular bulk collagen gels (Extended Data Fig. 3) or gels synthesized by other traditional approaches involving modulation of temperature^11^, pH^21^, or salt content^22^. Our collagen bundles can exceed 30μm in diameter. These μ-bundles have applications in tissue engineering, serving as mesoporous scaffolds with mesoscale topographical cues and enabling the study of cell-ECM interactions within more complex and physiologically relevant matrices containing thick and tortuous fiber bundles. By mixing thick collagen μ-bundles into hepatoblastoma cells (HepG2) during the multicellular spheroid formation (Extended Data Fig. 4), we create a set of pathophysiological liver tumor clusters, we term “fibrotic spheroids”, which highly resemble fibrosis regions in liver cancer patients (Extended Data Fig. 4b).

We next demonstrate our rapid collagen gelation technique for macroscopic patterning. We fabricate a polydimethylsiloxane (PDMS)-based multichannel device comprising parallel cylindrical channels (length: ∼ 5mm; diameter: 400μm) (Fig.1m, and fig.S5a,b). By flowing collagen solution through the channels, which are prefilled with PEG solution (Fig.1n, Movie 3), strip-like collagen gels of tunable lengths with high aspect ratio are molded by the channels and rapidly form within 2-3 minutes (Fig. 1o). We term these patterns “collagen strips”. The collagen strips can be easily manufactured using our device and subsequently collected or maneuvered. This molding technique enables a scalable approach for generating macroscopic collagen gels with high-aspect ratios. MMC-induced rapid gelation prevents cells from sedimenting during the course of gelation (Extended Data Fig. 5c-f), which can be utilized for fabricating multicellular tissues with improved cell distribution, which is demonstrated with human pluripotent stem cells (hPSCs) (Extended Data Fig. 5g,h) and liver cells (Extended Data Fig. 6).

Finally, the creation of complex, large-scale constructs can be accomplished by employing freeform bioprinting techniques, such as embedded 3D printing within PEG containing granular supporting baths (Fig. 1p). The collagen rapidly gels in place as the pattern is being extruded (Movie 4), resulting in rapidly formed and stable user-designable features (Fig. 1q, r).

### Vascular engineering with μ-bundles

Collagen μ-bundles are interactive with cells. Here, we explore the use of μ-bundles as tissue scaffolds for endothelial cells (ECs). Upon mixing ECs with μ-bundles, we observe that ECs physically recruit most μ-bundles within 1 hour and rapidly compact the resulting tissue within 6 hours (Extended Data Fig. 7a and Fig. 2a). The self-assembled EC-bundle construct is termed the ‘EC-patch,’ which contains a mesoporous microenvironment created by the μ-bundles. Over the course of 5 days, cells continue to compact the patch and migrate into its interior (Fig. 2b, c, and f). We generated EC-patches using different types of μ-bundles, distinguished as ‘Thick’ and ‘Thin’ bundles based on 10 mg/mL and 2 mg/mL collagen precursor solution, respectively (Extended Data Table 1). Notably, thick bundle patches exhibit significantly larger areas compared to thin bundle patches. Thick bundles gradually compact until reaching a plateau at Day 4, while thin bundle patches stabilize at Day 2, indicating that thick bundles are more resistant to cell contraction (Fig. 2c, Extended Data Fig. 7b). The thick μ-bundles-formed patches have a three-fold increase in surface area to volume ratio (Fig. 2d,e) that promoted cell viability (Extended Data Fig. 7c), infiltration (Fig. 2f), and interconnected EC network formation (Fig. 2h). ECs with proper junctions were identified by VE-Cadherin (Fig. 2g).

**Fig. 2.**
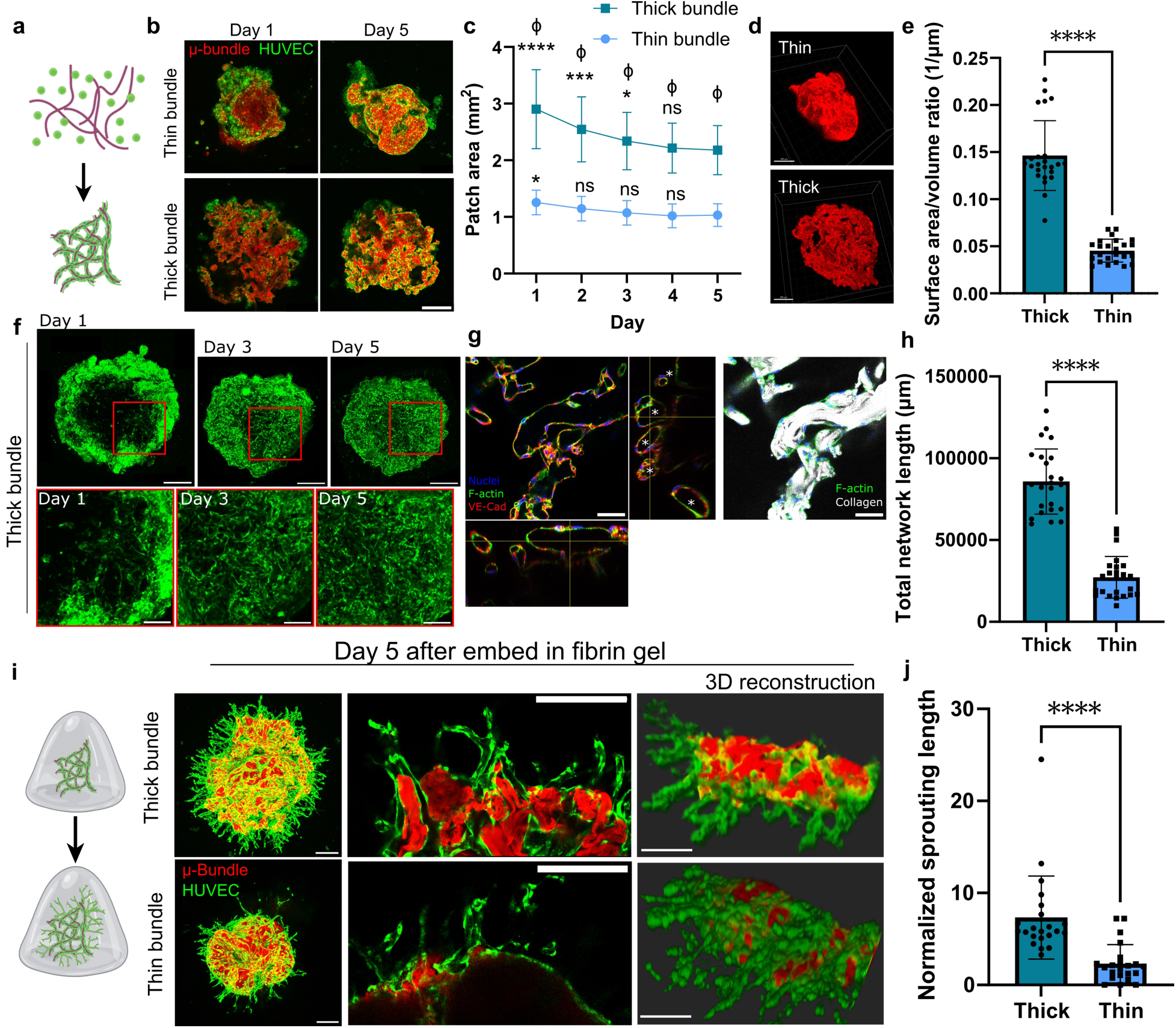
Fabricating functional EC-patches through self-assembly of cells and μ-bundles. (**a**) Schematic of EC-patch formation. HUVECs and bundles were loosely seeded, then self-assembled into a patch. (**b**) Live confocal images of patches with thick or thin bundles on Day 1 and Day 5, showing bundle guidance of the HUVECs. Images correspond to the max intensity projection of a z-stack around 200 μm. Scale bar, 500 μm. (**c**) Bundle patch compaction dynamics over 5 days. Statistical comparisons between “Thick” and “Thin” daily. * denotes comparing each day (Day 0-4) with Day 5 within groups. Ф denotes comparing two bundle groups on each day. ****P< 0.0001, ***P=0.0002. Thick bundle *P=0.0154, thin bundle *P=0.0204, determined by one-way ANOVA with Dunnett’s multiple comparisons (n=15 patches for each condition from N=3 biological replicates). Ф<0.0001, determined by unpaired t-test with two-tailed comparison. (**d**) 3D reconstruction of μ-bundle scaffolds after culturing for 5 days. Scale bars, 400 μm. (**e**) Quantification of surface area to volume ratio in the thick bundle patch and thin bundle patch, showing porosity of the structures (n=24; N=3). (**f**) Live confocal images showing daily tracking of patches demonstrating cell infiltration and network formation. Images correspond to the max intensity projection of a z-stack around 200 μm. Top, images showing an entire thick μ-bundle patch formation. Scale bars, 500 μm. Bottom, magnified images showing cell infiltration and network formation. Scale bars, 200 μm. (**g**) Fluorescent confocal images of representative patches on Day 5, VE-Cad (red), nuclei (blue), and actin filaments(green). μ-Bundles are labeled with Ester dye (gray). (**h**) Quantification of EC network length for each patch (n=24; N=3). (**i**) Patches were embedded in fibrin gel on Day 6 (schematic), showing the angiogenic sprouting ability of patches over 5 days. Top, patches with thick μ-bundle. Bottom, patches with thin μ-bundle. Scale bars, 300 μm. (**j**) Quantification of sprouting length of patches (n=24; N=3). All bar plots represent mean ± S.D. ****P < 0.0001, determined by a two-tailed student’s t-test.

To evaluate angiogenic potential, we embed the EC-patches into fibrin gels on Day 6 of the patch formation (Fig. 2i). Angiogenic sprouts with lumens emerge from the EC-patches, especially from thick bundle patches, and connect with the internal vascular network within the patches. Thick bundle EC-patches exhibit significantly higher sprouting length (Fig. 2j) and remain stable over 28 days (Extended Data Fig. 7d). These results demonstrated that EC-patches with thick bundles generate robust interconnected angiogenic sprouts with lumens (Extended Data Fig. 7e).

We further explore the function of the μ-bundles in the EC-containing macroscopic strips using TRACE method (Extended Data Fig. 8). Pre-formed thick μ-bundles can be mixed with collagen solution and ECs prior to generating tissue strips using the macroscopic strip method (Fig.1m-o and Extended Data Fig. 5-6). We show that the inclusion of bundles reinforces the tissue under cellular contraction (Extended Data Fig. 8a-d), provides guidance to ECs to form interior vascular network (Extended Data Fig. 8e-i), and promotes cell viability (Extended Data Fig. 8j). Overall, our results show that mesoscopic features conferred by μ-bundles in macroscopic tissue constructs facilitate lumenal and functional EC network formation.

### Deriving intestinal tubes from macroscopic strips

High aspect ratio tubular structures and strips can be found during development, such as neural tubes and gut tubes, and in the body, such as the gut tract and muscles. Our strip fabrication method enables scalable and rapid synthesis of macroscopic cell-laden strips with uniform cell distribution. We utilize this method to create macroscopic strip-like pluripotent tissues with hPSCs, which maintain pluripotency for at least a week (Extended Data Fig. 5g,h). Following established differentiation protocol^23^ for timing and media formulations that are commercially available (Fig. 3a), we differentiate the Day-1 pluripotent strips into intestinal tubes in basement membrane (BM) gels via *in situ* definitive endoderm (DE) induction for 3 days followed by the mid-hind gut induction for 5 days (Fig. 3b). The cells in the strip undergo cellular reorganization and express the DE marker SOX17 and the mid-hind gut marker CDX2. During the intestinal expansion, the strips evolve into self-organized tubular tissues with distinct proliferative (KI67 positive) crypt-like budding surrounding a central lumen - mimicking physiological intestinal tissue (Fig. 3c). The intestinal tissues are maintained for up to five weeks with increasingly more pronounced tissue budding (Fig. 3d). Mesenchymal cells are also generated around the intestinal buds and exhibit invasion into the local BM capsule (Fig. 3d). Moreover, our cultures achieve macroscopic scale and easily reach centimeter lengths (Fig. 3e). In Week 5 of the intestinal expansion, we observe the expression of villin and a small population of lysozyme positive cells (Fig. 3f,g), indicating the continuous maturation of the macroscopic tissues. These intestinal tubes can be fragmented and passaged, and the resulting fragments spontaneously form organoid clusters that can be further matured over a prolonged culture (Fig. 3h). Next, we expand the utility of the macroscopic strip by incorporating primary mouse intestinal cells. Comparing conventional, cluster-like intestinal organoids (Fig. 3i), we demonstrate that the macroscopic strips of isolated mouse intestinal crypts in BM encapsulation spontaneously assemble into centimeter-long, lumenal intestine-like constructs within 6 days. These primary intestinal tubes contain Paneth cells (lysozyme positive) rich crypt-like buds and β-catenin expressed at cell-cell junctions. In addition to the DE strips for intestinal organoids, we also show the hPSC strips can be induced into high aspect ratio embryoid bodies (EB) (Extended Data Fig. 5i) for downstream applications. We thus demonstrate a scalable method for generating macroscopic tissues derived from different cell sources in a physiological form.

**Fig. 3.**
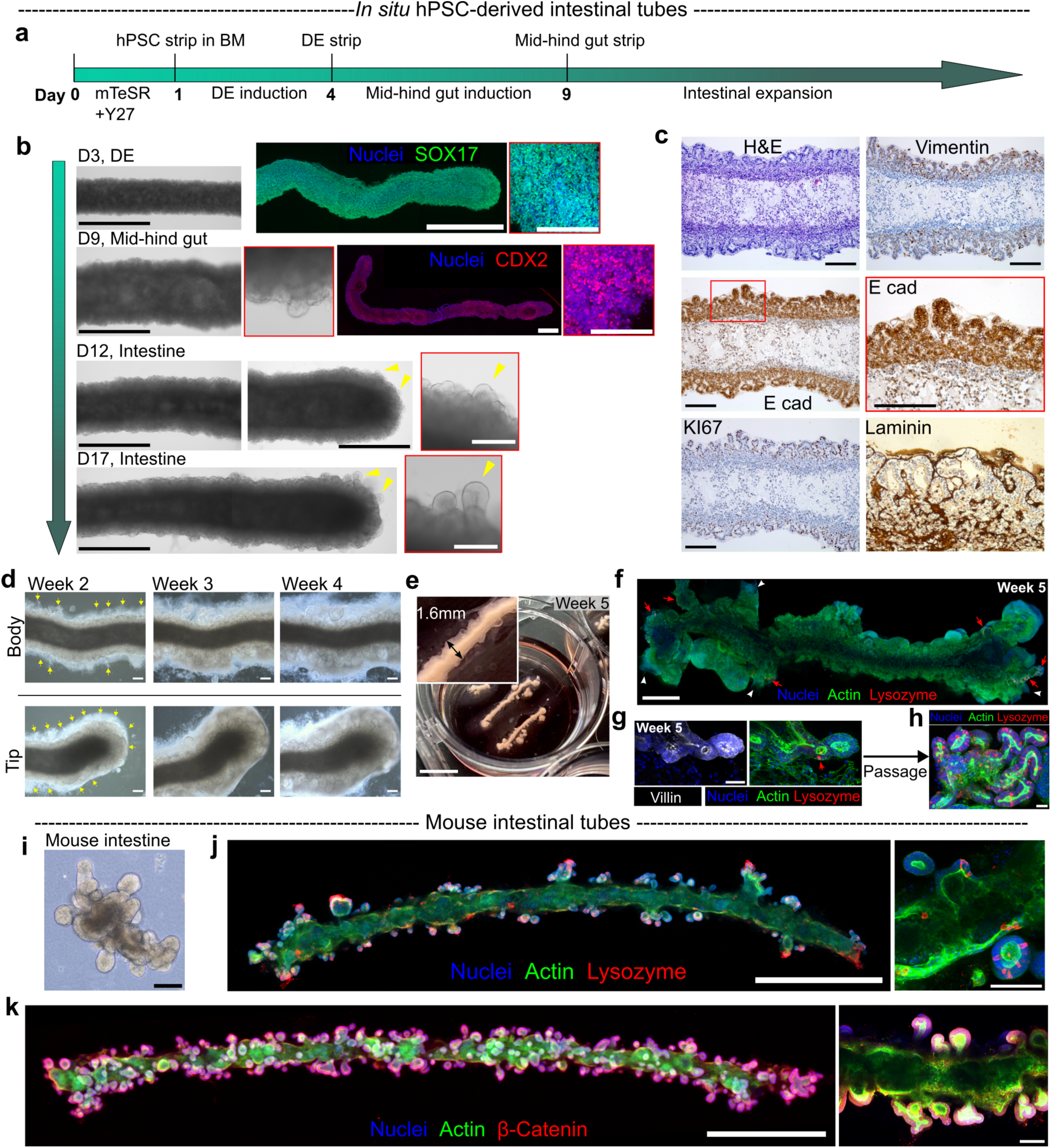
*In situ* differentiation and self-assembly of macroscopic intestinal tubes. (**a**) Timeline of differentiating the macroscopic hPSC strips into intestinal tubes *in situ*. (**b**) Representative brightfield images showing morphological changes of the strip and the cellular organization during the directed differentiation of the hPSC strip into an intestinal organoid over 17 days. Yellow arrowheads track the expansion of the same intestinal buds. The immunofluorescent staining confirms the differentiation stages: SOX17: DE marker; CDX2: mid-hind gut marker. Scale bars for large tissues: 1mm; scale bars for the close-up views, 200μm. (**c**) H&E and IHC staining on the intestinal tube after 20 days of differentiation from hPSC and 11 days of intestinal expansion show the hollow tube formation and spatially organized proliferative (KI67) intestinal buddings, the markers for premature intestinal organoids consisting of epithelial cells (E-Cad) and mesenchymal cells (vimentin), and cell polarity (laminin). Scale bars, 200 μm. (**d**) Long-term expansion (Week 2-4) of the intestinal tube leads to more pronounced intestinal budding formation and growth. Yellow arrows: mesenchymal cells. Scale bars, 200 μm. (**e**) Macrograph of centimeter-long, millimeter-wide hPSC-intestinal strips undergoing 5 weeks of expansion that exhibit pronounced buddings, scale bar, 10mm. (**f**) Immunofluorescent staining on a whole 5-week hPSC-intestinal strip showing cellular organization (actin) and intestine marker lysozyme (indicated with red arrows). Scale bar, 1mm. (**g**) Close-up imaging on the intestinal budding of a 5-week hPSC-intestinal strip showing the expression of intestine markers villin and lysozyme. Scale bar: 50μm. (**h**) Intestinal organoids passaged for 3 more weeks from the macroscopic 5-week intestinal strips show proper cell polarity and the enriched expression of lysozyme. Scale bar: 50μm. (**i**) Bright-field image of an organoid derived from the mouse small intestine. Scale bar, 100μm. (**j**) Fluorescent imaging of the mouse intestinal tubes self-assembled for 6 days exhibiting cell polarity and the tubular structure with buddings enriched with lysozyme. (**k**) Staining of the self-assembled mouse intestinal tubes showing the hollow tubular structure and membrane β-catenin. Scale bars in (**j**) and (**k**), 1mm. Scale bars in the close-up views: 100μm.

### TRACE-Bioprinting

Next, we present an enhanced embedded 3D collagen bioprinting based on the rapid gelation of collagen. To achieve this, we develop a support bath that mixes PEG solution with a shear-yielding agarose granular support bath, termed TRACE support bath (Extended Data Fig. 9a). While maintaining a shear-yielding behavior (Extended Data Fig. 9b,c), the TRACE support bath enables MMC during the freeform bioprinting process, which induces rapid gelation of both neutralized and acidic collagen at the desired location as the material is extruded from the print nozzle and comes into contact with the bath.

To demonstrate the printability of the collagen bioink within the TRACE bath, we utilize a commercial pneumatic bioprinter to extrude user-defined patterns in a layer-by-layer manner. We first print cell-free 3D structures with 6mg/mL acidic collagen ink in a free-standing manner within the support bath (Fig. 4a), such as thin-walled tubular geometries with various aspect ratios, hollow gourd shapes with spatially varying curvatures at different scales, multilayered “Y” logo, hollow stomach geometry, and pyramid with varied side profiles. The released objects closely matched the original computer-designed objects, indicating high fidelity for downstream tissue engineering applications.

**Fig. 4.**
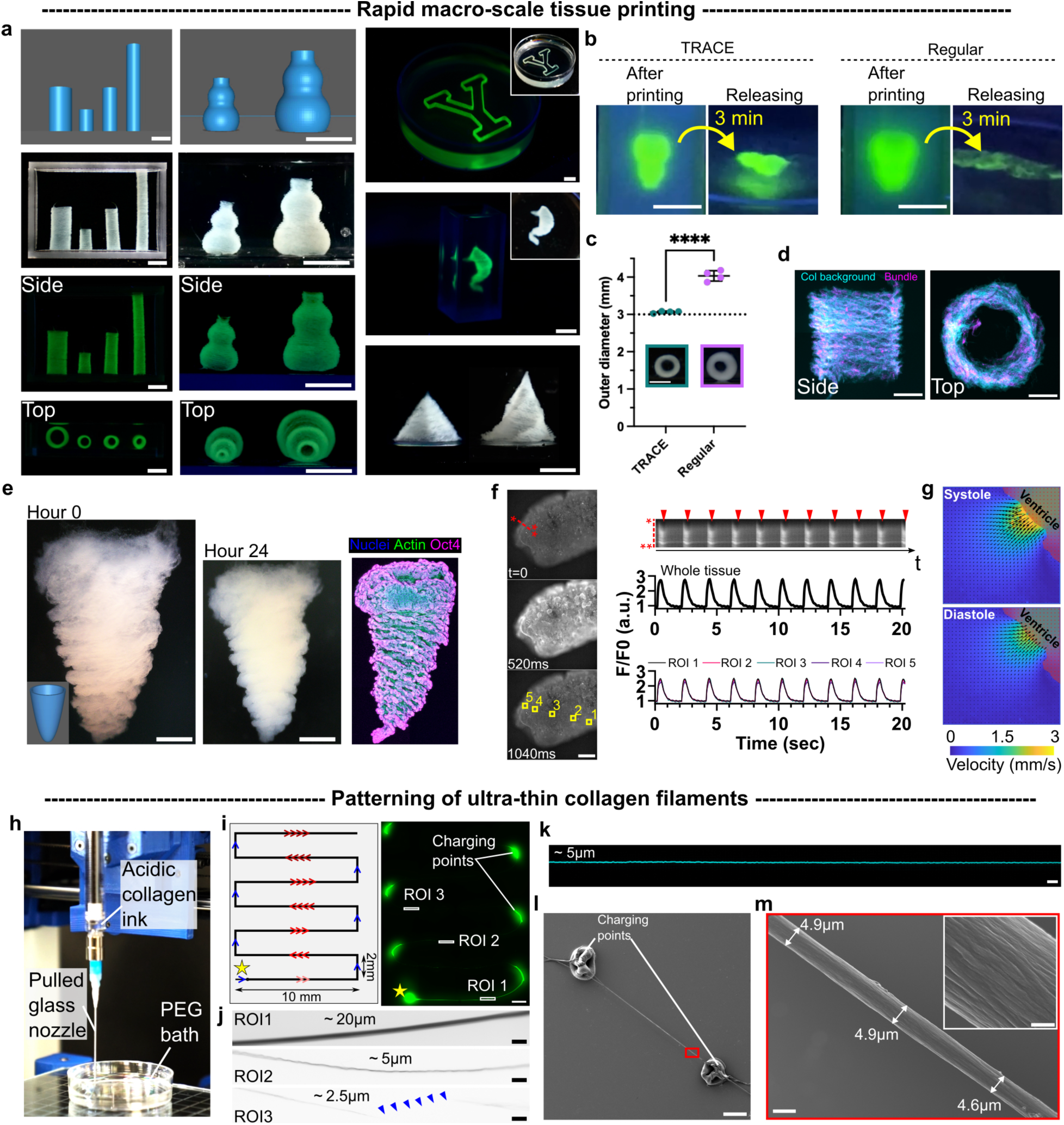
Multiscale rapid freeform printing of collagen. Enhanced macroscale 3D printing of cell-free or cell-laden collagen inks in TRACE support bath is demonstrated in (**a**) to (**g**). (**a**) Various macroscale constructs were 3D-printed using cell-free acidic 6mg/mL collagen. The first and second columns show the design and printed constructs released in PBS. The third column shows the printed structures in the support bath (fluorescently labeled) or stored in PBS. Scale bars, 5mm. (**b**) Releasing of 3D collagen constructs 3 min after printing in the TRACE vs. regular agarose support bath. Scale bar, 5mm. (**c**) Cross-sectional view of two tubes of neutralized 2mg/mL collagen printed in TRACE vs. regular bath and allowed to fully gel for 30 min. Scale bar, 3mm. A two-tailed student’s t-test was performed (****P<0.0001). (**d**) TRACE printed tubular structure with μ-bundle collagen composite ink. Scale bars, 1mm. (**e**) TRACE printed hPSC cardiac ventricle shell using neutralized 2mg/mL collagen ink) with tissue-relevant cell density (4×10^7^ cells/mL), and cultured for 24 hours. Scale bars, 1mm. Confocal imaging of the hPSC tissue after 24-hour growth stained with nuclei, F-actin, and Oct4. Scale bars, 1mm. (**f**) Characterizing TRACE-printed cardiac ventricle’s contraction and the synchronized calcium at both the whole-tissue level and the local level (ROIs 1-5). The kymograph indicates spontaneous contractions (red arrowheads) and calcium transient. Scale bar, 500 μm. (**g**) The cardiac cycle (systole and diastole) of the printed cardiac ventricle is indicated by PIV. (**h**) Bioprinting setup for ultrathin collagen filament patterning. (**i**) Fast printing of collagen (8mg/mL, acidic) in a raster pattern at various printing speeds (100mm/min, 3,000mm/min, 10,000mm/min, 12,000mm/min, indicated by varied numbers of red arrows), and with material “charging points’’ at a lower speed (70 mm/min, blue arrows). Star indicates the starting point of the printing path. (**j**) Regions of interest (ROIs) showing the widths of filaments printed at varying printing speeds. Blue arrowheads indicate the discontinuity of the filaments. (**k**) Confocal image of a single continuous ultrathin collagen filament printed with collagen labeled with GFP. (**l**) Scanning electron microscope of a patterned ultrathin collagen filament (∼5 μm thickness) created between two charging points. Scale bar, 200μm. The close-up image of the filament is shown in (**m**). Scale bar, 5μm. The surface topography of the single filament is shown in the inset. Scale bar in the inset, 1μm.

The improved effectiveness of the TRACE method for bioprinting was evaluated by printing the hollow gourd shapes in both the TRACE bath and a regular support bath (agarose slurry without the inclusion of PEG) (Fig. 4b, Movies. 5 and 6). The resulting structure was allowed to gel for 3 minutes before being released from the support baths. Notably, the TRACE-printed structure demonstrated MMC-induced rapid gelation and was easily released from the bath without disrupting its integrity. In contrast, the same design printed in the regular support bath did not maintain its structure and disintegrated after the releasing due to insufficient gelation, as unmodified collagen typically requires much longer gelation time (≥ 30 minutes)^24,25,17,18^.

We also demonstrated that MMC-induced gelation improved the printability of low-concentration unmodified collagen inks, which exhibit high fluidity and high diffusivity. We print tubular structures using a 2mg/mL neutralized collagen ink in either TRACE or regular support baths followed by the typical 30-minute gelation time (Extended Data Fig. 9d). Accelerated gelation by TRACE-printing shows improved accuracy with limited ink diffusion and reduced broadening of features (Fig. 4c, Extended Data Fig. 9e). In contrast, printing in the regular bath results in the inclusion of slurry material into the printed structures, porous collagen texture, and low fidelity (Extended Data Fig. 9f). These results highlight that MMC improves shape retention when printing unmodified low viscosity collagen. As demonstrated earlier (Fig. 2 and Extended Data Fig. 4), the collagen μ-bundles can guide cell migration, topography sensing, and EC network formation. To show the versatility of TRACE printing, we incorporated collagen μ-bundles (synthesized from the 2mg/mL precursor as described previously, Fig.1i,j) into a collagen ink and utilized this composite material to print a macroscopic tubular construct with tunable mesoscopic architectures (Fig. 4d).

To assess the biological performance of the low-concentration collagen bioink within the TRACE bath, 2D and 3D cell-laden structures are printed with neutralized 2mg/mL collagen bioink containing tissue-relevant cell densities. Specifically, HUVEC-containing ink (4×10^6^ cells/mL) is used to print 3D tubular and bell-shaped structures and a macroscopic 2D triangular pattern, followed by suspension culture of the printed tissues in the medium for 5 days (Extended Data Fig. 10a-d). Cell contractility is active in the printed tissues, inducing geometrically guided tissue compaction. Cells also form junctions and mimic vascular tissues, demonstrating physiologically relevant bioactivity (Extended Data Fig. 10e). Furthermore, using hPSC-containing collagen ink (4×10^7^ cells/mL) (Extended Data Fig. 10f) we print tubular tissue (Extended Data Fig. 10g) or cardiac ventricle-shaped tissues (Fig. 4e and Extended Data Fig. 10i). The bioactivity of the printed hPSC tissues is demonstrated by tissue compaction after 24-hour growth, maintenance of pluripotency markers OCT4 and Nanog (Fig. 4e, and Extended Data Fig. 10h), and self-organization (Extended Data Fig. 10j). We highlight that TRACE-enabled low concentration collagen printing supports pluripotent tissues, which lays the foundation for *in situ* differentiation of more complex tissue types, as we have demonstrated with the macroscopic strips (Fig. 4).

We further demonstrate TRACE-enabled functional tissue printing focusing on millimeter-scale cardiomyocyte ventricles. Prior methods of cardiac ventricle 3D bioprinting primarily rely on printing cell-laden fibrin bioink between bioprinted high concentration collagen layers^15^ or employing embedded printing with cell-laden gelatin microgel-based bioink^26^. Our approach offers a direct printing method using unmodified collagen-based ink loaded with hPSC-derived cardiomyocytes and human cardiac fibroblasts (HCFs). After 21 days of culture, the TRACE-printed cardiac ventricle exhibits whole-tissue spontaneous contractions, synchronized calcium transient (Fig. 4f, Movie 7), and fluid pumping cycles (Fig. 4g, Movie 8). These findings highlight the potential of TRACE-enabled functional printing in the application of cardiac tissue engineering.

### Pushing the limits of bioprinting resolution

We have demonstrated the fabrication of microscale collagen bundles through rapid mixing in a PEG solution. To achieve full controllability of the microscale collagen structures, we explore high-resolution bioprinting of collagen using TRACE methods with a customized fine printing nozzle (diameter: ∼ 20μm) (Fig. 4h, Extended Data Fig. 11a, and Movie 9). To induce instant gelation while creating a highly viscous bath without the agarose microgel support, we utilize a higher MW PEG (PEG20,000) at a higher concentration of 800mg/mL. In this PEG bath, the controlled nozzle rapidly pulls small amount of collagen into filaments from a pre-printed pool of collagen ink termed “charging points”. By printing raster patterns at varying printing speeds (Fig. 4i and j), we show that the nozzle speed is a key determinant of bioprinting resolution. At a high print speed (1,000-4,000 mm/min), we achieve high-resolution printing, generating continuous ultrathin collagen filaments measuring ∼ 5-10 μm in width. When printing speed reaches the order of 10^5^ mm/min, thinner filaments can be achieved but with occasional discontinuity. We extrude and pattern long, continuous filaments (Fig. 4k and Extended Data Fig. 11b-d), reaching macroscopic length scales while maintaining microscopic, single collagen fiber width^27,28^. Scanning electron microscopy (SEM) imaging (Fig. 4l,m) further reveals that the printed single collagen fibers are composed of highly packed and aligned fibrils, recapitulating hierarchically assembled collagen *in vivo*^29^. Thus, our method enables the bioprinting of macroscopic tissue patterns with microscopically resolved features.

## Discussions

In this study, we introduce a set of methods for patterning collagen using MMC, enabling highly controllable and advanced biofabrication of microscopic features and macroscopic collagenous tissues. The rapid gelation principle, induced by high MW PEG, facilitates the formation of collagen structures within seconds, allowing for diverse applications in engineering cell-matrix systems and bioprinting. Various basic geometries, including disks, microbeads, and strips, can be generated within minutes, while complex 3D collagen-based constructs can be printed and quickly gelled. This instant gelation method eliminates waiting time for gel formation, improving scalability and enabling more advanced tissue biophysics studies.

Recent advancements in bioprinting and organoids have led to the development of next-generation tissues and organoids that display physiologically relevant size, cellular organization, and functions^20,30–36^. Our macroscopic collagen patterns, densely packed with cells, facilitate the fabrication of realistic and functional tissue models, including complex geometries and macroscopic tissues from various cell types. These patterns support pluripotent stem cell maintenance and tissue differentiation in pre-defined configurations, making them valuable for preclinical drug screening and potential organ repair.

The TRACE-based μ-bundles mimic collagen architecture *in vivo*^37,38^. They are highly tunable with a wide range of bundle size, which elicits various cellular responses. The μ-bundles allow for the generation of microvascular networks and fibrotic regions within tumor spheroids, mimicking realistic liver tumors. Moreover, the high-resolution printing capabilities, down to subcellular scales, offer new possibilities for tissue and bioscaffold design, resembling the refined features observed in physiological tissues.

TRACE method provide enhanced bioprinting capabilities, enabling rapid and accurate printing of low viscosity, low concentration collagen bioinks. This advancement facilitates the creation of complex tissue models and supports various applications in tissue engineering, organoid development, and drug screening. The incorporation of microscale resolution in macroscopic collagen patterns opens up new avenues for tissue and bioscaffold design with improved fidelity and tunability.

## Methods

### Rapid gelation of collagen I in the macromolecular bath

The crowding bath was prepared by dissolving high molecular weight (8,000 and 20,000) polyethylene glycol PEG8000 (Sigma) and PEG20000 (Sigma) in sterile PBS at the concentration of 200mg/mL or 800mg/mL. The solution can be stored at room temperature. For tissue culture applications, the PEG bath was sterilized by a 0.45 µm syringe filter. Rat tail collagen type I with a stock concentration of 8-10mg/mL (Corning, 354249) was neutralized (pH =7.2∼7.4) with NaOH and adjusted to the desired concentrations with PBS. When delivered into the PEG8000 bath, a collagen solution rapidly polymerized and formed various shapes depending on the delivery methods.

### Fabrication of collagen disks via MMC

Macroscopic collagen disk was created by forming a small collagen droplet (tested volume: 1∼50 μL) at the tip of a micropipette tip, and then lowering the tip until the collagen droplet contacted and expanded the flat surface of the PEG bath. Each droplet instantly expanded and spread uniformly into a disk. This procedure can be repeated numerous times to render a large number of disks. These disks were washed at least three times in PBS. Circular microtissues were fabricated in the same way with a neutralized cell-laden collagen solution.

### Fabrication of collagen bundles via MMC

The μ-bundles were created by rapidly pipetting collagen solution (e.g.,200 μL) into a PEG bath (3 mL) with a 1000 μL pipette tip and quickly mixing for 20-30 times. To remove the PEG from the mixture before subsequent applications, 12mL PBS was added to the μ-bundles containing PEG bath and gently pipetted for a few times. The resulting suspensions were then centrifuged at 4000×g for 2 minutes, and a pellet of bundles was formed. The supernatant was discarded, and the bundle pellet was resuspended in 200 μL fresh PBS. The resulting μ-bundles could be incorporated into another hydrogel or mixed with cells to create a 3D tissue model with complex ECM architectures.

To study the effects of the concentration of the pre-bundle collagen solution on the bundle morphology, we used neutralized collagen with different concentrations (2, 4, 6, and 8mg/ml) and acidic collagen at 9.5-10 mg/ml. To conserve collagen concentration in the final solution, the resuspension volume was adjusted accordingly (Extended Data Table 1). For 2, 4, 6, and 8 mg/ml collagen μ-bundles, the initial neutralized collagen precursor was 200μL for each concentration. Then bundles were resuspended to 40, 80, 120, and 160 μL, respectively (Extended Data Table 1). Non-neutralized 9.5-10 mg/ml collagen μ-bundles were made with 200 μL stock collagen solution and resuspended to 200 μL. The washed PEG-free bundles were labeled by Alexa Fluor 647 NHS Ester (Succinimidyl Ester) (Invitrogen) diluted in PBS at the ratio of 1:1000 and incubated at 37°C for 30 minutes. The labeled collagen μ-bundles were washed in PBS, centrifuged, and resuspended again in previously described volumes. A Leica DMI8 confocal microscope was used to acquire images, and the diameter of bundles was quantified by an ImageJ plugin BoneJ ^39^.

### Fabrication of macroscopic collagen strips via MMC

Sterile PEG8000 solution (200mg/mL) was injected into the inlets of the PDMS multi-channel device. A neutralized collagen solution with or without cells was then gently injected into the PEG-filled channels. The collagen solution was allowed to stay in the channel for two more minutes for complete gelation into the cylindrical strips. The collagen strips were then carefully flushed out of the channels’ outlets with PBS via 200μL pipet tips from the inlets. These strips were collected in the Petri dish containing sterile PBS or fresh cell culture medium.

### Fabrication of collagen-based microparticles

With the same principle for making the disks, a large number of collagen microdroplets were sprayed into a PEG bath by plunging a syringe containing a small amount of collagen solution or by a paint spray airbrush. The collagen droplets polymerized into particles upon contact with the PEG bath.

### Collagen gel formation in the regular gelation conditions

Neutralized collagen solutions with different concentrations (2-9.5mg/mL) were prepared following the recipe stated in Extended Data Table 1. Droplets (2.5 μL) were dispensed on glass bottom dishes to form domes. For the acidic collagen stock (9.5mg/mL), 2.5 μL droplets of the solution were directly dispensed on the dishes without any modification. The collagen domes were quickly transferred to a humid incubator and allowed to gel for 40 minutes at 37℃.

### mMSC-laden collagen disks

The mMSC-laden disks were fabricated with 2mg/mL neutralized collagen solution with 4×10^5^ cells/mL. 30 μL cell-containing collagen droplets were dispensed on the PEG bath to form cell-laden disks for 3 minutes in a 100-mm Petri dish. The majority of the PEG bath was removed without disturbing the disks. 10-15mL PBS was added to rinse PEG off the disks. This procedure was repeated three times. MSC osteogenic induction medium (with or without 10 μM Y27632) was then added to replace the PBS. The media were changed every 3 days. The differentiation medium was the DMEM growth medium containing 50 μg mL−1 l-ascorbic acid (Sigma), and 10 mm β-glycerophosphate (Sigma). The osteogenic commitment was assessed by alkaline phosphatase (ALP) staining. ALP was stained using a FastBlue working solution of 500 μg/mL Fast Blue BB (Sigma) and 500 μg/mL naphthol-AS-MX (Sigma) phosphate in an alkaline buffer (100 mm Tris-HCl, 100 mm NaCl, 0.1% Tween-20, 50 mm MgCl_2_, pH = 8.2). On Day 9, the mMSC disks were fixed with 4% PFA and washed three times in PBS. The fixed samples were then equilibrated in the alkaline buffer for 15 minutes, followed by the incubation in FastBlue working solution for 60 minutes at room temperature. The samples were then washed in an alkaline buffer for 15 minutes, followed by 15 minutes in DPBS. ALP staining was imaged with a bright field microscope (Leica).

### Fabrication of the multichannel device for macroscopic collagen strips

The multichannel devices were fabricated using PDMS through a molding process. PDMS and silicone elastomer curing agent (SYLGARD^TM^ 184, Dow Corning, USA) were mixed in a 10:1 ratio and poured into a petri dish. Degassing in a vacuum chamber was performed to remove trapped air bubbles. To create parallel channels, 27 Gauge blunt needles (McMaster-Carr, IL, USA) were aligned and fully immersed in the PDMS liquid mixture. The mixture was subsequently cured overnight in the 60°C oven. After curing, the device was peeled off from the petri dish and trimmed into proper sizes, and the needles were demolded from the device. A 30-second treatment in a plasm cleaner (GLOW, Glow Research) was conducted on the device, to promote the channels’ surface hydrophilicity. For cell-laden strip fabrication, the device was sterilized with 70% ethanol, rinsed with sterile water, and air-dried in the tissue culture hood before use.

### Cell culture and maintenance

GFP-labeled human umbilical vein endothelial cells (Angio Proteomie, cAP-0001GFP) were cultured in 0.2% gelatin-coated T75 tissue culture flasks with EGM-2 medium (Lonza, CC-3162). The medium was changed every other day. Cells were lifted up by 0.25% Trypsin-EDTA (Gibco) once the confluency reached 70-80%. Then cells were passaged at the ratio of 1:3. Passage number 6 or lower was used for experiments. Human induced pluripotent stem cell (hPSC) line PGP1 was cultured with mTeSR1 (STEMCELL Technologies, #85850) in tissue culture 6-well plates coated with reduced growth factor Geltrex (Gibco, A1413202) diluted in cold DMEM/F12 (dilution: 1:30). The cells were passaged with ReLeSR (STEMCELL Technologies, #100-0484) when the confluency reached 80-90%, following the manufacturer’s protocol. The dissociated hPSC aggregates were allowed to attach in mTeSR1 containing 10μM Y27632 (Cayman) for 24 hours before maintenance in mTeSR1 with daily medium changes. Human hepatoma cell line HepG2 was cultured in T75 tissue culture flasks with complete DMEM containing 10% FBS and 1% Pen/Strep with a medium change every 2-3 days. The cells were passaged using 0.25% Trypsin-EDTA (Gibco) when the confluency reached ∼80%. Mouse bone marrow mesenchymal stem cells (BMSCs) (D1, ATCC CRL-12424) were cultured in T75 tissue culture flasks with complete DMEM containing 10% FBS and 1% Pen/Strep every 2-3 days. Passage number 12 or lower was used. Cells were collected once the confluency reached 70-80%. The cells were passaged at a ratio of 1:6 to 1:3 using Trypsin-EDTA. Human cardiac fibroblasts (HCFs, PromoCell, C-12375) were cultured in human cardiac fibroblast media (Cell Applications, 316-500), and media was changed every other day. Passage number 5 or lower was used.

### Generation of the EC-patch

96 well ultra-low adhesion plates (Corning, 3474) were used to generate and culture the EC-patches. Before seeding cells, bundles were adjusted to different volumes to conserve the 10 mg/mL concentration, as described in Extended Data Table 1. Bundles made from 2 mg/mL and 10 mg/mL collagen precursors were used. 100 μL HUVEC suspension with a density of 1×10^6^ cells/mL was added to each well and then mixed with 5 μL bundle suspension in each well, rendering approximately 1×10^5^ cells and a total 50 μg collagen mass per patch. 10 mg/mL bundles were prone to be entangled and clumped after the centrifugation and re-suspension procedures. To ensure a more homogenous bundle suspension and thus a uniform incorporation with the cells, surgery scissors were applied to chop the original bundles into shorter and discrete sizes. A Leica DMI8 confocal microscope was used to acquire images. Images were taken at Hours 0, 1, 6, and 24. After 24 hours, images were taken every day. The EC-patches were cultured for 5 days, and the medium was changed every day.

### Angiogenesis assay for EC-patches

The EC-patches cultured for 6 days were embedded in 3mg/mL fibrin gels for the angiogenesis assay. In brief, 15 mg fibrinogen (Sigma, F8630) was dissolved with 2.5 mL PBS in a 37 ℃ water bath for at least three hours and sterile-filtered by a 0.22 μm PVDF syringe filter (Millipore, SLGVR33RS). The concentration of the fibrinogen stock solution was 6 mg/mL. Each EC-patch was transferred by a wide bore pipette tip into 50 μL EGM-2 medium containing 4 U/mL thrombin (Sigma, T9549). Then the medium with a patch was mixed with 50 μL fibrinogen stock solution. A 100 μL fibrin gel dome with the patch was plated in a well of a 24-well plate. This procedure was repeated to make multiple gel domes encapsulating an EC-patch. These fibrin gel domes were then incubated in a 37 ℃ incubator for 15 min. After the fibrin gels were fully solidified, 1 mL EGM-2 medium supplemented with an additional 75 ng/mL VEGF (Gibco, PHC9394) was added to each well. The medium was changed every other day. Confocal live imaging was taken every day for one week. The long-term culture of EC-patches in the fibrin gels was up to 28 days.

### Fabrication of the EC-strip

A 4 mg/mL neutralized collagen solution was prepared on ice. The prepared collagen was mixed with the same volume of cell suspension with a density of 8×10^6^ cells/mL. The final density of HUVECs and the collagen concentration of the mixture were 4×10^6^ cells/mL and 2 mg/mL, respectively. For the strips with bundle gels, 100 μL 4 mg/ml neutralized collagen precursor with HUVECs (8×10^6^ cells/mL) was made. Then collagen precursors with cells were mixed with 100 μL 10 mg/mL bundle solution homogenized with surgical scissors. The EC-strips with or without bundles were rapidly fabricated with the multichannel device. In brief, HUVEC/bundle-containing collagen precursors were pipetted into the PEG8000-filled PDMS channel. The strips were allowed to fully gel for 2 minutes before being washed out with sterile PBS. The strips were transferred and cultured in 24-well ultra-low attachment plates (Corning, 3474) with EGM-2 medium for 5 days. 1 mL EGM-2 medium was added to each well with a medium change every other day. Strips were moved to glass bottom plates (Cellvis) before imaging, and a confocal microscope (Leica DMI8) was used to take images every day.

### Isolation and maintenance of mouse intestinal crypts

The Mouse IntestiCult medium (STEMCELL Technologies) was prepared and supplemented with 100 U/mL Pen/Strep (Gibco), according to the manufacturer’s manual. A C57BL/6 mouse was pre-sacrificed under the approval of Yale University animal protocol (YD000223). A small intestine (∼20 cm long) near the stomach was harvested in ice-cold PBS supplemented with Pen/Strep. The intestine segment was washed by cold PBS to flush out the content and cut open along the axial direction. The intestine was further cut into 2-mm small pieces with sterilized surgical scissors. These segments were transferred to 15 mL fresh cold PBS in a 50 mL conical tube and rinsed three times. The segments were then incubated in 25 mL Gentle Cell Dissociation Reagent (STEMCELL Technologies) at room temperature for 15 minutes on a rocking platform at 20 RPM. After the tissue segments were settled, the supernatant was discarded, and 10 mL of cold PBS with 0.1% BSA was added. The tissues were then mixed by pipetting up and down. The supernatant was filtered through a 70 μm cell strainer and collected in a new 50 mL conical tube. This step was repeated twice. Next, the filtrate was centrifuged at 4 °C, 290 *g* for 5 minutes. The supernatant was discarded, and the pellet containing intestinal crypts was resuspended in 10 mL cold PBS with 0.1% BSA and centrifuged at 4 °C, 200 *g* for 5 minutes. The supernatant was discarded, and the crypts were resuspended in 10 mL, 4°C DMEM/F12 (Gibco). Crypts were centrifuged at 4°C for 5 minutes again and resuspended again with 300 μL DMEM/F12 medium. The crypt suspension was mixed with 300 μL growth factor reduced Matrigel (Corning, 354230) or growth factor reduced Geltrex (Gibco, A1413202) and dispensed as gel domes in 24-well tissue culture plates, followed by 15-minute gelation in the 37°C incubator. The crypt-containing gel domes were cultured in the IntestiCult medium with a medium change every two days. The intestinal organoids grown from the crypts were passaged after 1 week. Domes were treated by Gentle Cell Dissociation Reagent for 1 minute and were broken by pipetting up and down. Suspension was transferred to 15 mL conical tubes and centrifuged at 4 °C, 290 *g* for 5 minutes. Then the supernatant was discarded. Pellet was resuspended in 10 mL cold DMEM/F12. The suspension was then centrifuged again at 4 °C, 200 *g* for 5 minutes. The supernatant was gently removed. The pellet was resuspended in IntestiCult medium, and the same volume of Matrigel was added. Matrigel domes were seeded in 24 well tissue culture plates and 50 μL for each dome. The medium was changed every two days.

### Fabrication of macroscopic mouse intestinal tubes

Mouse intestinal organoids were collected before experiments. 1 mL gentle cell dissociation reagent (STEMCELL Technologies) was added to each well. Then Geltrex domes were broken by pipetting up and down. The organoids from 20 domes were collected in a conical tube and treated in the gentle cell dissociation reagent for 7 minutes. Then the organoids suspension was centrifuged at 4°C, 290 *g* for 5 minutes. The supernatant was discarded, and 10 mL 4°C DMEM/F12 was added to resuspend the pellet. The suspension was centrifuged again at 4°C, 200 *g* for 5 minutes, the supernatant was gently discarded. The pellet was then resuspended in 1mL of neutralized 2 mg/mL collagen and centrifuged at 4°C, 200 *g* for 5 minutes to enrich the fragmented organoids at the bottom. Then excessive collagen solution was removed. The PDMS channels were filled with sterile 200 mg/mL PEG 8000, and 6 μL concentrated organoid bioink was pipetted into each channel. After 2 minutes, intestinal strips were flushed out by sterile PBS into a PBS bath. Then the strips were collected and embedded in Geltrex and cultured in 6 well tissue culture plates for 6 days. 3 mL IntestiCult medium was added to each well, and the medium was changed every 2 days.

### Patterning of ultrathin collagen microfilament

An open-source extrusion-based 3D printer (Lulzbot) was used to pattern the microscale fibers. A pulled glass nozzle (inner diameter: 15 μm, WPI) was coaxially connected to the Gauge 25 needle and tightly sealed with Parafilm. Acetic collagen (8-10 mg/mL) was delivered into the fine glass nozzle via the interior needle at designed extrusion speeds. For the microfiber patterning, millimeter-scale 2D patterns were designed with SolidWorks (Dassault Systèmes) and converted the dxf files to G-Codes with a free online G-Code generator (https://optlasers.com/cnc-software/g-code-generator). The G-Code was then modified to be compatible with the 3D printer (Lulzbot), and print filaments at a wide range of speeds (100mm/min, 3,000mm/min, 10,000mm/min, 12,000mm/min). The nozzle was allowed to proceed at a lower speed (70mm/min or 0.3mm/min) at each turning point to build excessive material defined as a “material charging point. The G-Codes for different patterns with designated extrusion speed and printing speed were included in the supplementary information.

### *In situ* derivation of the intestinal tubes from hPSC-strips

On Day 0, ∼80% confluent human induced pluripotent stem cells (hPSCs) were dissociated into single cells with the Gentle Cell Dissociation Reagent (STEMCELL Technologies). Bioink consisting of 1.5mg/mL collagen with 5% Geltrex and high-density hPSCs (4×10^7^ cells/mL) was prepared by mixing equal volumes of 3mg/mL neutralized collagen with 10% Geltrex and 8×10^7^ cells/mL cell suspension in mTeSR 1(STEMCELL Technologies) containing 10μM ROCK inhibitor Y27632 (Cayman). The bioink was rapidly crowded in the multi-channel device with a 400μm channel diameter to generate the macroscopic tissue strips. These hPSC strips were subsequently rinsed in a PBS bath and then maintained at 37°C in mTeSR 1 with Y27 for several hours. Individual strips with a small amount of medium were then carefully moved to pre-chilled micro-centrifuge tubes and then mixed with cold reduced growth factor basement membrane matrix Geltrex (Thermo Fisher). The dilution of Geltrex by the introduced medium was less than 20%. A wide bore pipet tip was used to pick up ∼100 μL Geltrex solution with a single hPSC strip and slowly lay out the strip in an elongated gel dome onto a petri dish surface. After 20-minute gelation at room temperature, the hPSC strips in the Geltrex were covered with Y27 containing mTeSR and cultured overnight. On Day 1, the Geltrex domes with the hPSC strips were gently detached from the petri dish and cultured in fresh mTeSR without Y27 for 6 hours to promote cell self-assembly. Once the strips showed smooth boundaries, as an indication of proper cell merging and polarity, we initiated the intestinal organoid differentiation on the strips using a commercially available kit STEMdiff™ Intestinal Organoid Kit (05140, STEMCELL Technologies) containing the necessary medium for each differentiation stage. We strictly followed the manufacturer’s technical manual to prepare the inductions media. With the DE induction medium, the hPSC strips were first differentiated into DE strips for 3 days with daily medium change. To guarantee a full encapsulation in the basement membrane niche, the floating DE strips were embedded once again in high-density Geltrex that was anchored on a 6-well plate. With the mid-hind gut induction medium, these embedded DE strips were then induced into mid-hind gut strips for 5 days with daily medium change. In the intestinal Organoid Growth Medium (OGM) with the OGM supplement and 1% GlutaMax (Thermo Fisher), the mid-hind gut strips were allowed to develop and mature in the Geltrex capsule for up to 5 weeks into intestinal-specific tubes.

### Induction of EB strips

High cell density (4×10^7^ cells/mL) hPSC strips containing 1.5mg/mL collagen with 5% Geltrex were collected in an Anti-Adherence Rinsing Solution (STEMCELL Technologies) treated 60mm Petri dish. These hPSC strips were suspended and cultured in mTeSR medium with 10μM Y27632 for 24 hours. At Hour 24, the mTeSR medium was then gently removed and replaced by the EB Formation Medium (STEMCELL Technologies). The strips were cultured in the EB Formation Medium for 5 days with daily medium change.

### Subculture of the hPSC-derived intestinal tubes into matured intestinal organoids

The hPSC-derived intestinal tubes expanding for 5 weeks were extracted from the Geltrex capsule with ice-cold DMEM/F12 (Thermo Fisher) and broken down to small fragments by a 200uL pipet tip rinsed with Anti-Adherence Rinsing Solution (07010, STEMCELL Technologies). The tissue fragment suspension was transferred into an Anti-Adherence Rinsing Solution coated 15mL conical tube on ice and allowed to sediment by gravity for at least 5 minutes. The tissue fragments were purified after clearing out dead cell debris by gently removing the supernatant, resuspending in 2mL fresh DMEM/F12, and centrifuge-pelleting at 300 RPM. The purified tissue fragments with the minimum medium were suspended and plated in 40-50 μL Geltrex per 3D dome in each well of a 24-well plate. After full gelation for 30 minutes at room temperature, the Geltrex domes were covered by 500 μL Intestinal OGM with 1% GlutaMax per well. The intestinal tube fragments were cultured with a full medium change every 2-3 days, and passaged with the same subculture procedure every 1-2 weeks.

### Cardiac differentiation from hPSCs

hPSCs were maintained in mTeSR1 until they had above 98% confluency before starting differentiation. Differentiation to cardiomyocytes was performed as previously described^40,41^ and summarized in Extended Data Table 2. In brief, on day 0 of differentiation, we prepared 12.5 μM CHIR99021 (STEMCELL Technologies, 99021) in a mixed medium of mTeSR1 and RPMI/B-27 without insulin (v/v was 1:3); RPMI (Gibco, 11875-093) and B-27 without insulin (Gibco, A18956-01) were mixed beforehand at a ratio of 49:1. 24 hours later, media was replaced with RPMI/B-27 without insulin. On Day 3 at the same hour when CHIR99021 was added, the medium was replaced with RPMI/B27 without insulin, containing 5 μM IWP4 (STEMCELL Technologies, 72552). An additional 48 hours later (on Day 5), medium was replaced with RPMI/B27 without insulin, and media was refreshed on day 7. On day 9, media was replaced with RPMI/B27 (Gibco, 17504-044), and media was refreshed on day 11. On day 12, we purified cardiomyocytes using RPMI without glucose, containing 4 mM lactate (L1750-10G)^42^, and media was refreshed on Day 14. On Day 16, media was replaced with RPMI+B27. On Day 18, cardiomyocytes were dissociated using TrypLE (Gibco, A1217702) following its commercial protocol.

### Histology on macroscopic tissue strips

The HepG2 strips and the hPSC-derived intestinal strips in the Geltrex capsules were fixed in PBS solution with 4% PFA overnight at 4°C, and embedded in melted HistoGel (Epredia) in a plastic cryomold. The tissue-containing HistoGel was allowed to solidify at room temperature, demolded, and preserved in 4% PFA at 4°C. The samples were sent to Yale Pathology Tissue Services (YPTS) for paraffin embedding, sectioning, and staining. The embedded tissues were sectioned with a thickness of 10 μm. Hematoxylin Eosin (H&E) staining were conducted for both tissue types. The HepG2 strips were also stained with PAS (Periodic acid-Schiff’s reaction) and PAS after treatment with diastase to remove glycogen (D.PAS). Using immunohistochemistry (IHC), the hPSC-derived intestinal tubes were stained for KI-67, vimentin, E-cadherin, and Laminin.

### TRACE support bath generation with Agarose granular gel/PEG mixture

Biocompatible inert hydrogel, agarose, was utilized to create a supportive slurry following a protocol modified from a previous study^43^. In brief, the agarose slurry was prepared by dissolving 0.7%(w/w) agarose (Sigma-Aldrich, MO, USA) in boiling 50mL PBS (Gibco, NY, USA) in a 100mL glass media bottle, followed by gradual cooling from 85 °C to 20 °C on heat plate while stirring with a magnetic bar at 700 rpm. TRACE support bath was created by thoroughly mixing 5 parts of the resulting agarose slurry with 1 part of 800mg/mL PEG 8000 solution in PBS (Gibco, NY, USA) to obtain a final PEG concentration of 200mg/mL. Mixing was achieved by pumping the two materials in two 10mL syringes (BD Emerald, NY, USA) back and forth through a 90-degree syringe luer lock connector (McMaster-Carr, IL, USA) for 100 times. For cell-laden collagen bioprinting, the 0.7% agarose solution was autoclaved prior to cooling, and PEG 8000 was filtered through a 0.45mm filter (Millipore) for sterilization. The mixing accessories were also autoclaved. For cell-laden macroscale tissue strip assembly, cell culture medium-rich granular support was created by mixing the culture medium with an agarose granular support bath in a 1:1 ratio.

### TRACE bioink preparation

Three bioinks were employed in TRACE 3D printing studies. To evaluate the printability of TRACE methods with low-concentration collagen, acellular bioink was prepared in two concentrations. The 6mg/mL acidic collagen was prepared by diluting the stock collagen at a concentration of 8.08mg/mL (354249, Corning, NY, USA) with pre-chilled 0.02N acetic acid (Fisher, IL, USA). The 2mg/mL neutralized collagen was prepared following the method described in the previous section. Food color (Market Pantry) and 1-μm yellow-green fluorescent beads (F8823, Invitrogen) were incorporated into the acellular bioink to aid the visualization of the printed construct. To prepare the bioink of cell-laden collagen, neutralized collagen at a concentration of 2mg/mL was made following the previously described method. HUVECs and hPSCs were then mixed with neutralized collagen to produce a final concentration of 4 × 10^6^ and 4 × 10^7^ cells/mL, respectively. The bioink for ultrathin collagen filament printing was the acidified stock collagen (8-9.5mg/mL) or the stock collagen mixed with 10% type I collagen conjugated with Alexa-Fluor 647 for fluorescent imaging. For the printed cardiac tissue, bioink was prepared by combining dissociated cardiomyocytes and HCFs in a neutralized 3mg/mL collagen with 15% Geltrex (v/v) with cell concentrations of 4×10^7^ cells/ml for hPSC-derived cardiomyocytes and 4×10^6^ cells/ml for HCFs, following a previously established protocol^44^.

### TRACE printing

Extrusion-based bioprinting was performed using a commercialized pneumatic bioprinter Cellink BioX6 (CELLINK, Gothenburg, Sweden) with the temperature-controlled print head. All digital models were designed in SolidWorks (Dassault Systemes SolidWorks Corp, Velizy, France) and exported as stereolithography (.stl) files. In general, 3D structures were designed with diameters ranging from 3mm to 6mm, and heights ranging from 3mm to 20mm. The files were sliced and printed using DNA Studio 3.0 (Cellink, Gothenburg, Sweden), with a layer height of 0.2mm. 25-gauge nozzles with 0.5 or 1 inch in length (0.5 or 1inch) (McMaster-Carr, IL, USA) were utilized to print at speeds of 10mm/s or 5mm/s, with extrusion pressures of 4-8kPa for acellular printing and 20kPa for cell-laden printing. Prior to printing, a container with sufficient size to accommodate the printed construct was filled with the TRACE support bath and secured to the printing bed. The temperature-controlled print head was set to 8°C. Each ink was loaded into a pre-chilled cartridge (CELLINK, Sweden) through a straight socket-to-socket connector (51525K423, McMaster-Carr, IL, USA) and then stored on ice until use. To ensure sterile conditions during cell-laden collagen printing, the printer chamber was subjected to a built-in UV cycle, and the clean chamber fan equipped with a HEPA filter was activated. The cartridges and nozzles used for printing were sterilized by autoclaving. After the printing was completed, the printed structures were secured in the TRACE bath for at least 3 minutes, and then released carefully from the bath with a spatula or by directly pouring. The printed construct was then thoroughly rinsed in a PBS bath to remove any excess TRACE bath. Subsequently, the printed construct was transferred to desired culture media and cultured in a humid incubator at 37°C with 5% CO_2_ for future applications.

### Maintenance of cardiac pumping tissues

Following a previously established protocol^41^, the bioprinted cardiac tissues were cultured in seeding media (10% FBS (Gibco, 10438-026), 1% P/S (Gibco, 15140122), 1% NEAA (Gibco, 11140-050), 1% L-glutamine (Gibco, 25030-081), and 1% sodium pyruvate (Gibco, 11360-070), and 10 μMY27632 (Stemcell Technologies, 129830-38-2)) on day 0. On day 1, media was refreshed with DMEM (Gibco, 11995065)/B27(Gibco, 17504-044) mixed at a ratio of 49:1 (v/v), and media was refreshed every other day from then on.

### Calcium imaging

We followed a previous protocol on calcium imaging^15^. Tissues were first incubated at 37°C for 90 minutes in Tyrode’s solution containing 5 µM calcium indicator Cal 520 AM (AAT Bioquest, 21130) and 0.025% Pluronic F127 (Sigma Aldrich, P2443). Tissues were then incubated in Tyrode’s solution for 30 minutes at 37°C for equilibration. Finally, media was replaced with DMEM/B27 during imaging in an environmental chamber at 37°C, 5% CO_2_.

### Particle imaging velocimetry (PIV) measurements

Particle imaging velocimetry (PIV) measurements were performed using a Leica DMi8 system at 10x magnification with a mercury lamp and a high sensitivity Hamamatsu camera. Tissue culture medium consisted of 0.5% (v/v) 1 μm-diameter fluorescent microspheres (ThermoFisher Scientific, # F8821) to visualize fluid flow. The time interval of imaging was set to 17 milliseconds. PIV analysis was performed using PIVlab ^45^ in MATLAB (MathWorks). Imaging was conducted in DMEM/B27 and in an environmental chamber at 37°C, 5% CO_2_.

### Rheometric characterizations

Gelation kinetics of collagen at room temperature or the rapid collagen gelation triggered by MMC were characterized on an Anton Parr rheometer (502 WESP). The top plate was a 25-mm round glass coverslip (VWR) and the bottom plate was a 40-mm round coverslip (Fisherbrand). To prevent slip, both plates were functionalized by polydopamine coating. The rheometer was then zeroed and calibrated. Immediately after, approximately 350mL of a neutralized collagen solution (2mg/mL) was dispensed between the plates with a 500μm gap. The sample was kept in a custom humidity chamber to prevent evaporation. Gelation was monitored by imposing a 0.5% strain at a frequency of 0.2 Hz. The shear modulus of the collagen was monitored at room temperature for 200 seconds. For the rapid gelation, after the collagen solution was dispensed and while the time sweep started monitoring, PEG8000 solution (200mg/mL in PBS) was added to the sides of the parallel plates. For the rapid gelation, after the collagen solution was dispensed and while the time sweep started monitoring, PEG8000 solution (200mg/mL in PBS) was added from the sides of the parallel plates. The total length of the time sweep was 200 seconds. Mechanical properties (shear modulus) of the agarose slurry bath and the TRACE printing bath were characterized on a TA Instruments rheometer (Discovery HR 30), using a stainless steel 40.0mm, 1.98833° cone plate at room temperature. Both strain sweep (0.01%-100%) at the frequency of 1Hz and frequency sweep (0.1-100 Hz) at the strain of 0.1% were performed on the shear rheometer.

### Immunofluorescent staining

The EC-patches and EC-strips were fixed on Day 5 with 4% Paraformaldehyde (PFA, Santa Cruz Biotechnology) for 10 minutes and then washed with PBS three times. 1% BSA in PBS was used to block the samples at room temperature for 1 hour. The samples were incubated with primary antibodies overnight at 4°C. The samples were then washed with PBS for 3 hours and incubated with Hoechst (1:2000), rhodamine phalloidin(1:1000, Abcam), and a secondary antibody overnight at 4°C. The samples were washed with PBS for 3 hours on the next day before imaging. The macroscopic hPSC constructs, hPSC-derived intestinal tubes, and mouse intestinal organoid tubes were fixed in 4% PFA at room temperature for 30 minutes and washed with PBS three times. Samples were permeabilized by 0.2% Triton-X100 (Sigma) for 1 hour at room temperature and then washed with PBS three times. Samples were then blocked by 1% BSA overnight at 4°C. These samples were then incubated with primary antibodies overnight at 4°C, followed by washing with PBS for at least 3 hours at room temperature. The samples were incubated at 4°C overnight with secondary antibodies, Hoechst (1:2000), and rhodamine phalloidin (Abcam, 1:1000). The samples were washed with PBS for at least 3 hours before imaging.

### Cell viability assay

To evaluate cell viability of the GFP-labeled HUVEC patches and the macroscopic HUVEC strips, propidium iodide (PI)-based Cell Viability Kit (R37610, Invitrogen) was used to stain dead cells. On Day 5 of the experiments, the cellular constructs were incubated with the staining medium containing PI at the ratio of 2 drops per 1 mL medium. After 25-minute incubation at 37°C, the cells were imaged at 535/617 nm on a confocal microscope to detect the total GFP-labeled cells and the dead cells. For HepG2 strips, the LIVE/DEAD Cell Imaging Kit (R37601) was used to label live (green) and dead (red) cells. After 25-minute incubation at room temperature, the strips were imaged at 488/570 nm excitation on a confocal microscope. Viability was calculated as a percentage of PI stain-negative cells out of the total cells.

### Scanning electron microscopy

Patterned ultrathin filaments were collected by the more visible charging points with tweezers and rapidly transferred into fresh PBS. After a thorough rinse three times, the filaments were fixed in 2.5% glutaraldehyde (Polysciences, Inc.) diluted in deionized water for 20 minutes. The filaments were then rinsed with deionized water and collected on a clean glass coverslip. The collected filaments were allowed to be air-dried at room temperature overnight. The coverslip with the filaments was then mounted on a support with carbon tape (3M) and then coated with a 4nm layer of iridium with a sputter coater. A Hitachi SU-70 scanning electron microscope (magnifications: 50×, 2,000×, and 10,000×) was used to image the printed filaments with an accelerating voltage of 5 kV and an emission current of 29000 nA.

### Characterizations on the HUVEC-bundle patches

#### Surface area to volume ratio of collagen networks

Imaris 10.0 (Oxford Instruments, UK) was used to 3D-reconstruct the compacted bundles and extract the bundle structure’s surface area and total volume on each EC-patch. For the 3D reconstruction, we set the surface detail to 4.55μm and the absolute intensity threshold to 53.4. The same setting was applied to measure all the patches. The ratio was computed by dividing the surface area (*S_patch_*) by the total volume (*V_patch_*):

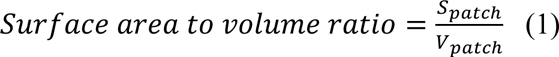

#### Total network length of the EC-patch

Maximum intensity projection (MIP) was applied to the confocal image stack of each EC-patch. The ImageJ plugin AngioTool ^46^ was then used to extract the total networks. For each patch, the extracted network was skeletonized, and the total length of the skeleton was measured.

#### Normalized sprouting length of the angiogenesis assay

MIP of the confocal images was applied to each EC-patch embedded in fibrin gels. The angiogenic sprouting of an EC-patch was considered as the total network grown from the boundary of the patch into the surrounding fibrin gel, defined as *netL_patch_*, extracted by AngioTool. The extracted network was skeletonized, and the total length of the skeleton was measured. The perimeter of the pre-angiogenic patch (*P_patch_*) was manually measured. The normalized sprouting length for EC-patches was computed by dividing the sprouting network length (*netL_patch_*) by the patch perimeter (*P_patch_*):

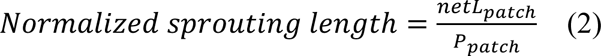

### Characterizations on the HUVEC-strips

EC networks in the strips were evaluated by normalized network length. The normalized network length was computed by dividing the measured total network length (*netL_strip_*) per strip by the axial cross-sectional area (*S_strip_*) per strip:

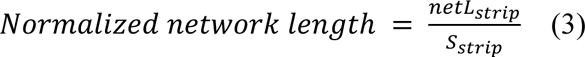

### Statistical analysis

Sample sizes and P values are reported in the figure legends and statistical analyses were processed in GraphPad Prism 9. Statistical differences between sets of data and analytical methods are reported in the figure legends. The difference is considered significant when the P value is less than 0.05. Comparisons with no significant difference are labeled as “ns”. The calculated P values are stated either in the figure or in the figure legends if they are greater than 0.0001. Otherwise, the convention for P values is: ****P < 0.0001 or #P < 0.0001. All bar plots represent mean ± standard deviation (SD), unless otherwise stated in the figure legends. Boxes, lines in the boxes, and the whiskers in the box plots represent 25th to 75th percentiles, median, minimum, and maximum. Three lines of the violin plots indicate the median, the 25th, and 75th percentiles.

### Antibodies

Primary antibodies: anti-VE-Cad (2158S, Cell Signaling Technology, 1:200), anti-Lysozyme (PA129680, Invitrogen, 1:50), anti-Villin (sc-58897, Santa Cruz, 1:50), anti-β Catenin (D10A8, Cell Signaling Technology, 1:50), anti-SOX17 (D1T8M, Cell Signaling Technology, 1:50), anti-CDX2 (D11D10, Cell Signaling Technology, 1:50), anti-Nanog (sc-293121, Santa Cruz, 1:50), anti-OCT4 (#2750, Cell Signaling Technology, 1:200). Secondary antibodies: Alexa Fluor 647 Goat Anti-Mouse IgG antibody (Invitrogen), and Alexa Fluor 488 Goat Anti-Rabbit IgG antibody (Invitrogen), Alexa Fluor 647 Goat Anti-Rabbit IgG antibody (Invitrogen).

## Acknowledgment

We thank Michelle Wu, Dr. Catherine Kim, Dr. Anjelica Gonzalez for the access to the CELLINK bioprinter and the training on the equipment. We also thank Zewu Zhu and Dr. Clemens Bergwitz for providing a dissected small intestine from a sacrificed mouse. We thank Dr. Stuart Campbell’s lab, Xia Li, and Ilhan Gokhan for the hPSC line PGP1 and the technical assistance in cardiac differentiation. We acknowledge the fundings: Yale Liver Center Pilot Project under award NIH P30 DK034989 (M.M.), NIH R35GM142875 (M.M.), and NIH grant T32EB019941 (R.Y.N).

## Author contributions

X.G. and M.M. conceptualized and designed this study. X.G., Z.W. and S.L. performed multiscale patterning of collagen and EC-patch experiments. Z.W. and S.L. performed experiments on EC-patch and angiogenesis. Z.W. performed isolation of mouse small intestinal cells and organoid maintenance. X.G. and Z.W. performed *in situ* hPSC differentiation and macroscopic mouse intestinal tubes. X.G. and Z.L. designed the TRACE support bath and performed the bio-printing of collagen. H.X. conducted cardiac differentiation, maintenance, imaging, and the developed the collagen-cardiomyocytes bioink. X.G., Z.W., Z.L. and S.L. analyzed and visualized data. X.G. and T.W. designed and fabricated the PDMS multichannel devices. X.G. and R.Y.N. performed rheological characterization. X.G. and R.Y.N. prepared SEM samples and performed SEM. A.R. helped with bioprinting of ultrathin filaments and visualization of μ-bundle formation. M.M. supervised the study. X.G., Z.W., Z.L. and M.M. wrote the manuscript with all the authors’ input.

## Data availability

All data supporting the results of this study are available within the Article and its Extended Data. All raw data are available from the corresponding author upon reasonable request.

## Competing interests

Authors declare that they have no competing interests.

## Notes

### Competing Interest Statement

The authors have declared no competing interest.

