## Supplementary Materials for "Instant Assembly of Collagen for Scaffolding, Tissue Engineering, and Bioprinting"

**Supplementary Text**
**Extended Data Figs. 1 to 11**
**Extended Data Tables 1 to 2**
**Movies 1 to 10**

### Supplementary Text

#### Rapid patterning of cell-rich, functional tissues

Our collagen patterning methods can be used for versatile cell patterning and tissue engineering applications, including at high, tissue-relevant cell densities. To create cell-laden gels, we can use pre-neutralized collagen solutions, reducing cell stress caused by high acidity. We can also use relatively low collagen concentrations (e.g. 2mg/mL), enabling sufficiently large porosity to promote cell spreading, cell-cell interactions, and self-organization. High collagen concentrations can lead to exceedingly small pore sizes that reduce cell motility. Here, we demonstrate several basic examples of engineered tissues created through rapid gelation of cell-rich collagen solutions.

Mesoscopic bundle networks can be used for tissue scaffolding (Extended Data Fig. 4a). By mixing thick collagen μ-bundles into hepatoblastoma cells (HepG2) during the multicellular spheroid formation, we create a set of pathophysiological liver tumor clusters, we term "fibrotic spheroids". In the fibrotic spheroids, we observe not only cell-cell interactions but also cell-collagen interactions, which morphologically resemble the fibrotic regions in liver cancer patients (Extended Data Fig. 4b). As seen in patient histology samples of fibrotic liver tumors, the tumor configuration often contains densely packed tumor cells with fibrotic inclusions in the tumor interior. Furthermore, these fibrotic spheroids exhibit larger sizes compared to the control spheroids made without collagen or with only neutralized collagen solution (Extended Data Fig. 4c,d). This demonstration provides a novel tumor model that potentially better recapitulates fibrosis conditions in cancer for high-throughput drug screening.

Next, we generate high aspect ratio macroscopic strips containing high-density liver cells (Extended Data Fig. 6) or human induced pluripotent stem cells (hPSC) (Extended Data Fig. 5). Compared to conventional thermal gelation (Extended Data Fig. 5c,d), MMC-induced rapid gelation of low concentration collagen prevents cells from sedimenting during the course of gelation, thus rendering a much more uniform cell distribution (Extended Data Fig. 5e,f). By rapidly molding a pre-neutralized collagen-cell mixture using our microchannel technique (Extended Data Fig. 6a), we form cohesive and densely packed liver strips (Extended Data Fig. 6d), which display high viability (Extended Data Fig. 6b) and liver metabolic activities, such as glycogen storage labeled by periodic acid-Schiff (PAS) staining (Extended Data Fig. 6c). Meanwhile, the hPSC strips can expand for a long period of time while maintaining pluripotency (Extended Data Fig. 5g,h). Inspired by the previous works[30, 47, 48], we further demonstrate the rapid molding technique provides a simple and fast way to fabricate tissue building blocks with a tunable aspect ratio for large-scale complex tissue assemblies (Extended Data Fig. 6e).

Using our droplet/disk method, we generate collagen disks filled with mouse mesenchymal stem cells (mMSCs) (Extended Data Fig. 2a). The mMSCs compact the gels over time, and the Rho kinase (ROCK) inhibitor Y27632 reduces the compaction by inhibiting the actomyosin contractility (Extended Data Fig. 2b). During the gel compaction, the cells also spontaneously fold the disks from the outer region (Extended Data Fig. 2c,d) and differentiate into an osteogenic phenotype under osteogenic induction (Extended Data Fig. 2e). This demonstrates the applicability of our method for cell mechanics (e.g. ECM compaction), tissue morphogenesis (e.g. macroscale tissue folding), and stem cell differentiation studies. Each of our patterning techniques (Fig.1) is compatible with both cell-free and cell-rich tissue engineering applications. For cell-rich applications, cells and a neutralized collagen solution can be pre-mixed together, creating a cell-laden ink.

**G-code for ultrathin collagen filament printing**

**Ultrathin filament printing: Raster lines with increasing speed:**

```
M221 S100 T0
G92 E0
M107

; Origin is current position
G92 X0 Y0 Z0
;=============================================================
;=============================================================
G1 X1 F30 E0.02
G1 X10 F100 E0.04
G1 X10 Y2 F70 E0.06
G1 X0 Y2 F3000 E0.08
G1 X0 Y4 F70 E0.10
G1 X10 Y4 F3000 E0.12
G1 X10 Y6 F70 E0.14
G1 X0 Y6 F10000 E0.16
G1 X0 Y8 F70 E0.18
G1 X10 Y8 F10000 E0.20
G1 X10 Y10 F70 E0.22
G1 X0 Y10 F12000 E0.24
G1 X0 Y12 F70 E0.26
G1 X10 Y12 F12000 E0.28

G0 Z30 F500
G0 X0  F1000
```

**Ultrathin filament printing: Windmill pattern**

```
M82 ;absolute extrusion mode
M221 S100
M73 P0 ; clear GLCD progress bar
M75 ; Start Print Timer

M221 S100 T0
G92 E0
M107

; Origin is current position
G92 X0 Y0 Z0

;=============================================================
G1 X0 Y0.01 F0.3 E0.03 ;origin
```

```
G1 X0 Y9.99 F4000 E0.06
G1 X0 Y10 F0.5 E0.09

G1 X4.99 Y8.65 F4000 E0.12
G1 X5 Y8.66 F0.5 E0.15

G1 X0.01 Y0.01 F4000 E0.18
G1 X0 Y0 F0.5 E0.21

G1 X8.65 Y4.99 F4000 E0.22
G1 X8.66 Y5 F0.5 E0.23

G1 X9.99 Y0 F4000 E0.23
G1 X10 Y0 F0.5 E0.24

G1 X0.01 Y0 F4000 E0.24
G1 X0 Y0 F0.5 E0.27

G1 X8.65 Y-4.99 F4000 E0.27
G1 X8.66 Y-5 F0.5 E0.28

G1 X4.99 Y-8.65 F4000 E0.28
G1 X5 Y-8.66 F0.5 E0.29

G1 X0.01 Y-0.01 F4000 E0.29
G1 X0 Y0 F0.5 E0.32

G1 X0 Y-9.99 F4000 E0.31
G1 X0 Y-10 F0.5 E0.32

G1 X-4.99 Y-8.65 F4000 E0.33
G1 X-5 Y-8.66 F0.5 E0.34

G1 X-0.01 Y-0.01 F4000 E0.35
G1 X0 Y0 F0.5 E0.38

G1 X-8.65 Y-4.99 F4000 E0.39
G1 X-8.66 Y-5 F0.5 E0.40

G1 X-9.99 Y-0.01 F4000 E0.41
G1 X-10 Y0 F0.5 E0.42

G1 X-0.01 Y0 F4000 E0.43
G1 X0 Y0 F0.5 E0.46
```

```
G1 X-8.65 Y4.99 F4000 E0.47
G1 X-8.66 Y5 F0.5 E0.48

G1 X-4.99 Y8.65 F4000 E0.49
G1 X-5 Y8.66 F0.5 E0.50

G1 X-0.01 Y0.01 F4000 E0.51
G1 X0 Y0 F0.5 E0.53


G0 Z30 F500
G0 X0  F1000
```

**Ultrathin filament printing: Spiral pattern**

```
M82 ;absolute extrusion mode
M221 S100
M73 P0 ; clear GLCD progress bar
M75 ; Start Print Timer

M221 S100 T0
G92 E0
M107

; Origin is current position
G92 X0 Y0 Z0

G0 X-0.5 F0.5 E0.03
G3 X-12.5000 I-6.0000 J0.0000 F3000 E0.06
G3 X-2.5000 I5.0000 J0.0000 F3000 E0.09
G3 X-10.5000 I-4.0000 J0.0000 F3000 E0.12
G3 X-4.5000 I3.0000 J0.0000 F3000 E0.15
G3 X-8.4163 I-1.9581 J0.0000 F3000 E0.18
G3 X-6.5001 I0.9581 J0.0000 F3000 E0.21

G0 Z30 F500
G0 X0  F1000
```

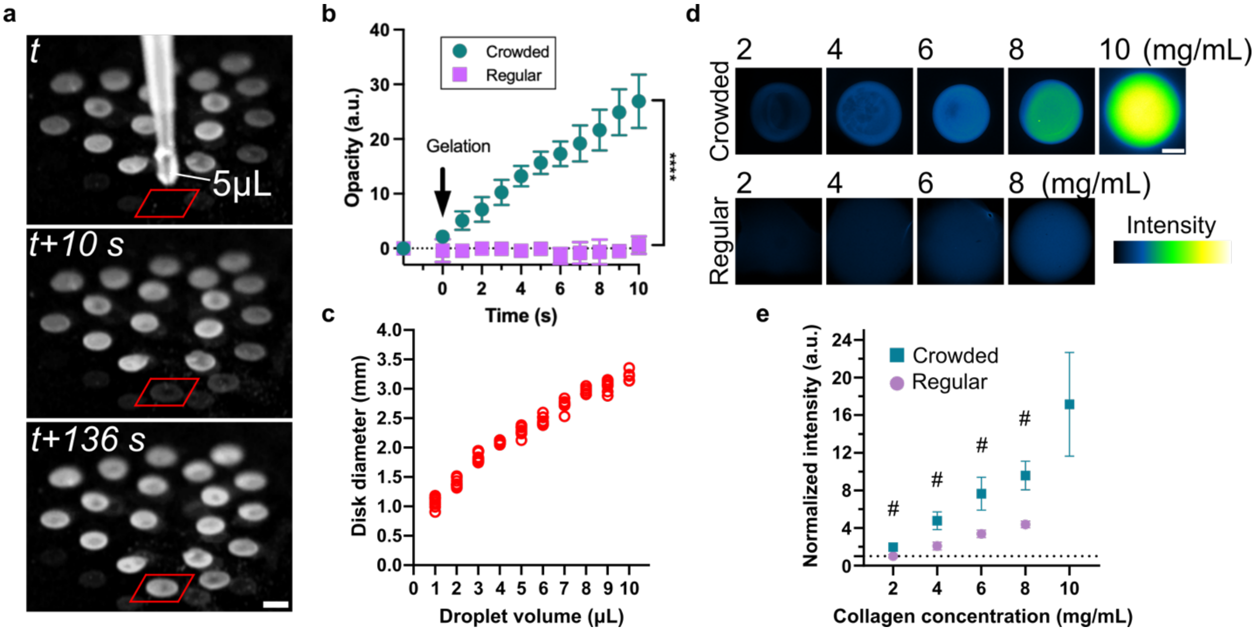


**Extended Data Fig. 1. High throughput fabrication of μ-liter disks.** (a) Video recording of μ-liter collagen disk fabrication in a high throughput way. The red box indicates the location of the PEG bath surface where a 5 μL collagen droplet was deposited at t=0. The appearance of the same collagen gel was shown at t=10s and t=136s. Scale bar, 2mm. (b) Opacity change for the first 10 seconds after the collagen droplets were dispensed in the PEG bath or the PBS bath. The disks showed opacity within 1 second and became increasingly visible, while the droplets in PBS did not appear and showed no opacity against the black background. Comparisons were made between the opacities of two conditions using a two-tailed student's t-test (****$P<0.0001$). (c) Disk diameter quantification as a function of the volume of the collagen precursor droplet, demonstrating the robust fabrication of the disks with tunable size. (d) Confocal images showing the fluorescent intensity of crowded disks and regularly gelled collagen domes made of GFP-labeled collagen with varied collagen concentration. Scale bar: 500μm. (e) The fluorescent intensity measurements of the gels normalized against a regularly gelled 2mg/mL collagen dome. Comparisons were made between crowded disks and regular neutralized bulk gels at different collagen concentrations using a two-tailed student's t-test (#$P<0.0001$). This shows that MMC enhanced the overall density of the collagen gels.

------------------------------ Mesenchymal stem cell disks --------------------------

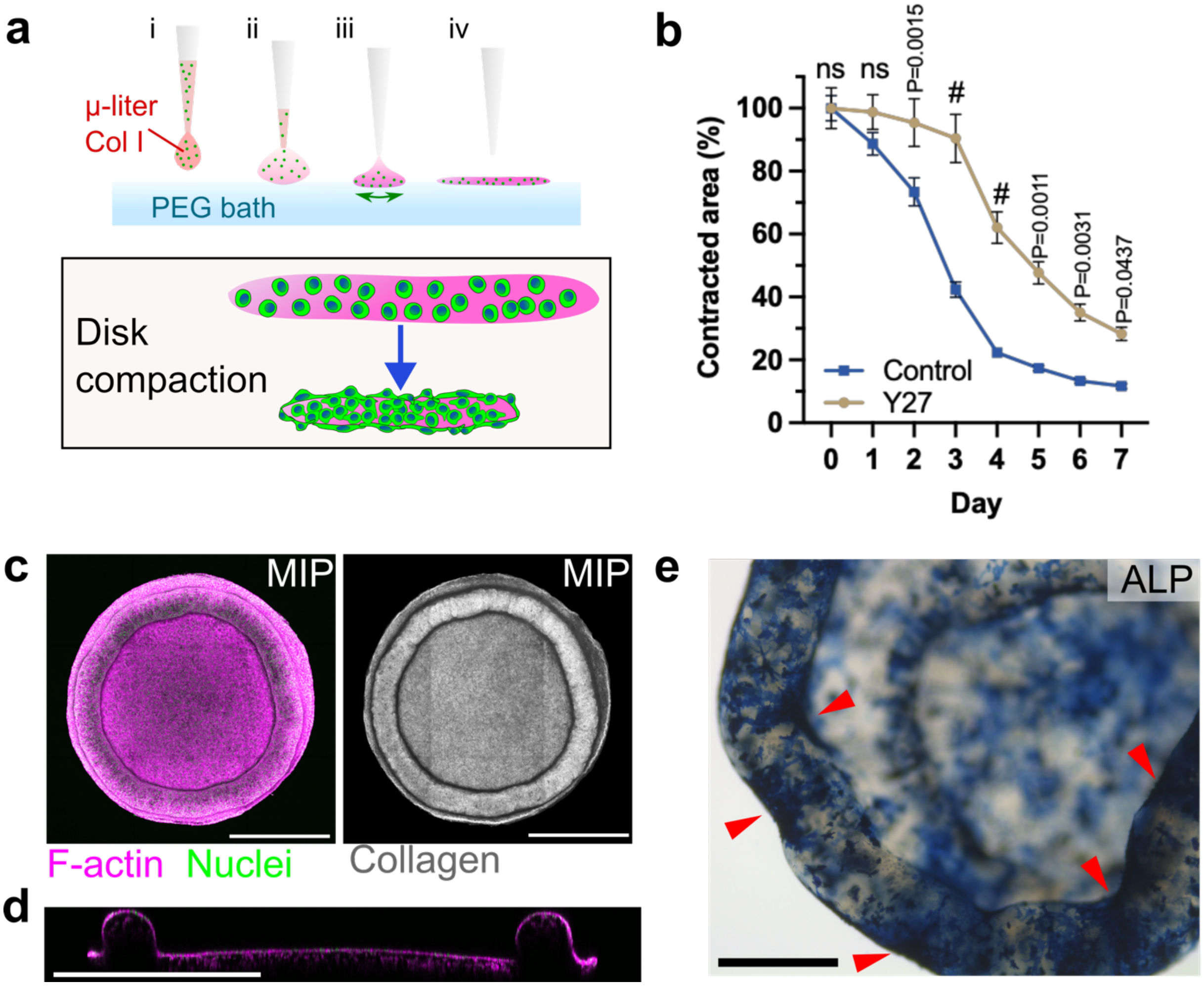


**Extended Data Fig. 2. Cell-laden collagen disks fabricated using TRACE method.** (a) Schematic of producing cell-laden μ-liter collagen disks to study cell contraction-induced collagen compaction. (b) Tracking compaction of mMSC-laden μ-liter disks with and without ROCK inhibitor Y27632 treatment for 7 days. #P<0.0001, determined by two-way ANOVA with Šídák's multiple comparisons (n=16-20 disks from N=2 independent replicates each condition). Data is represented as mean± SEM. (c) Max intensity projection (MIP) of a confocal fluorescent image stack of a Day-7 mMSC disk (F-actin and nuclei) and global tissue folding induced by cellular force (reflectance mode: collagen). The cross-section profile along the diameter of cells on the folded disk is shown in (d). Scale bars in (c) and (d), 1mm. (e) Staining of alkaline phosphatase (ALP) on the Day-9 mMSC disks. Red arrows indicate regions enriched with osteogenic mMSCs. Scale bar, 500 μm.

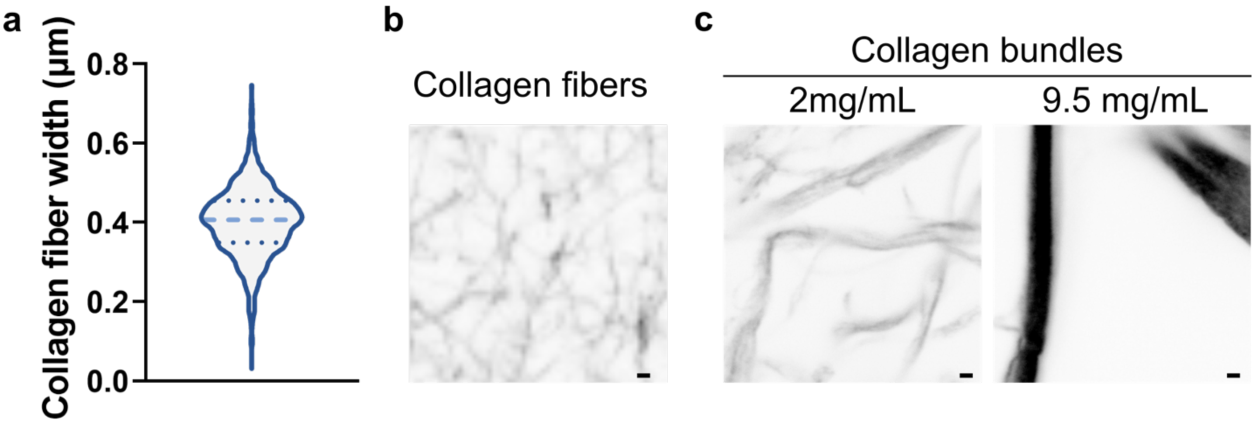


**Extended Data Fig. 3. Comparing collagen nanofibers and TRACE collagen bundles**. (a) Fiber width quantification of 2mg/mL regular collagen gels (gelled at 37°C for 40min without PEG). The gels were fluorescently labeled and imaged with a confocal microscope. The measurements were pooled from three fields of view from three independent collagen gels. The fibers have sub-micron (nanoscale) widths. (b) Representative image of collagen fibers. (c) Representative images of collagen bundles made with 2mg/mL and 9.5mg/mL pre-bundle collagen solutions. Scale bars, 1µm.

---------------------------- Fibrotic liver spheroids --------------------------------

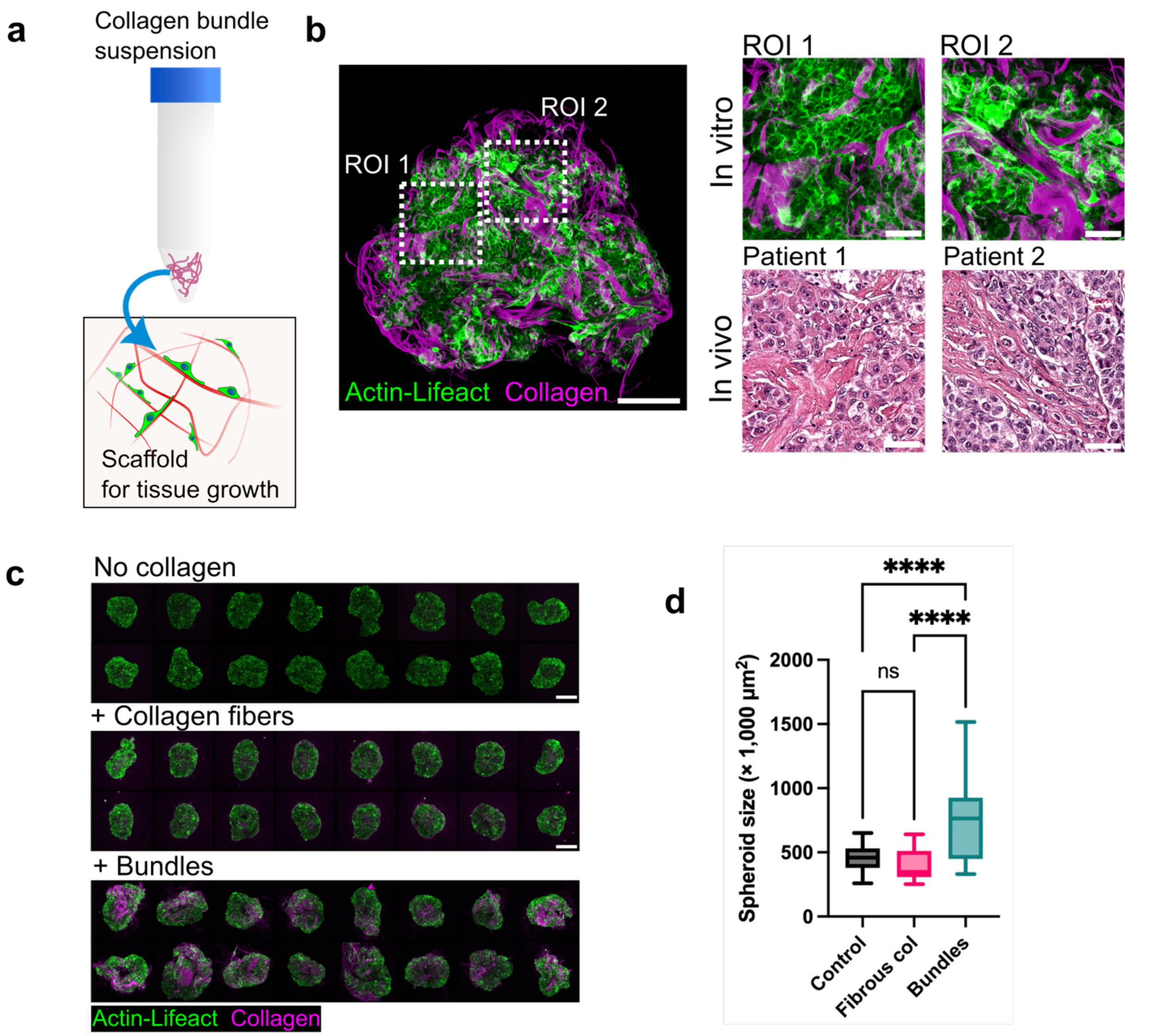


**Extended Data Fig. 4. Multicellular tissue with mescoscopic collagen architectures.** (a) Schematic of using collagen μ-bundles as a novel ECM scaffold for cell culture. (b) High-resolution imaging of a fibrotic liver spheroid Scale bar, 200 μm. Bundle and cell morphology in ROI1 and ROI2 are compared with those in local fibrotic regions of liver cancer patients. Scale bars in ROIs and histology slides, 50 μm. (c) High-throughput production of three types of LifeAct-tagged liver cell HepG2 spheroids: (i) without collagen, (ii) mixed with the neutralized collagen solution during formation, and (iii) mixed with thick collagen μ-bundles. Scale bars, 400 μm. (d) Size measurements of the three types of spheroids shown in (b). ****$P < 0.0001$, determined by one-way ANOVA with Tukey multiple comparisons (n=32 spheroids each condition from N= 2 replicates).

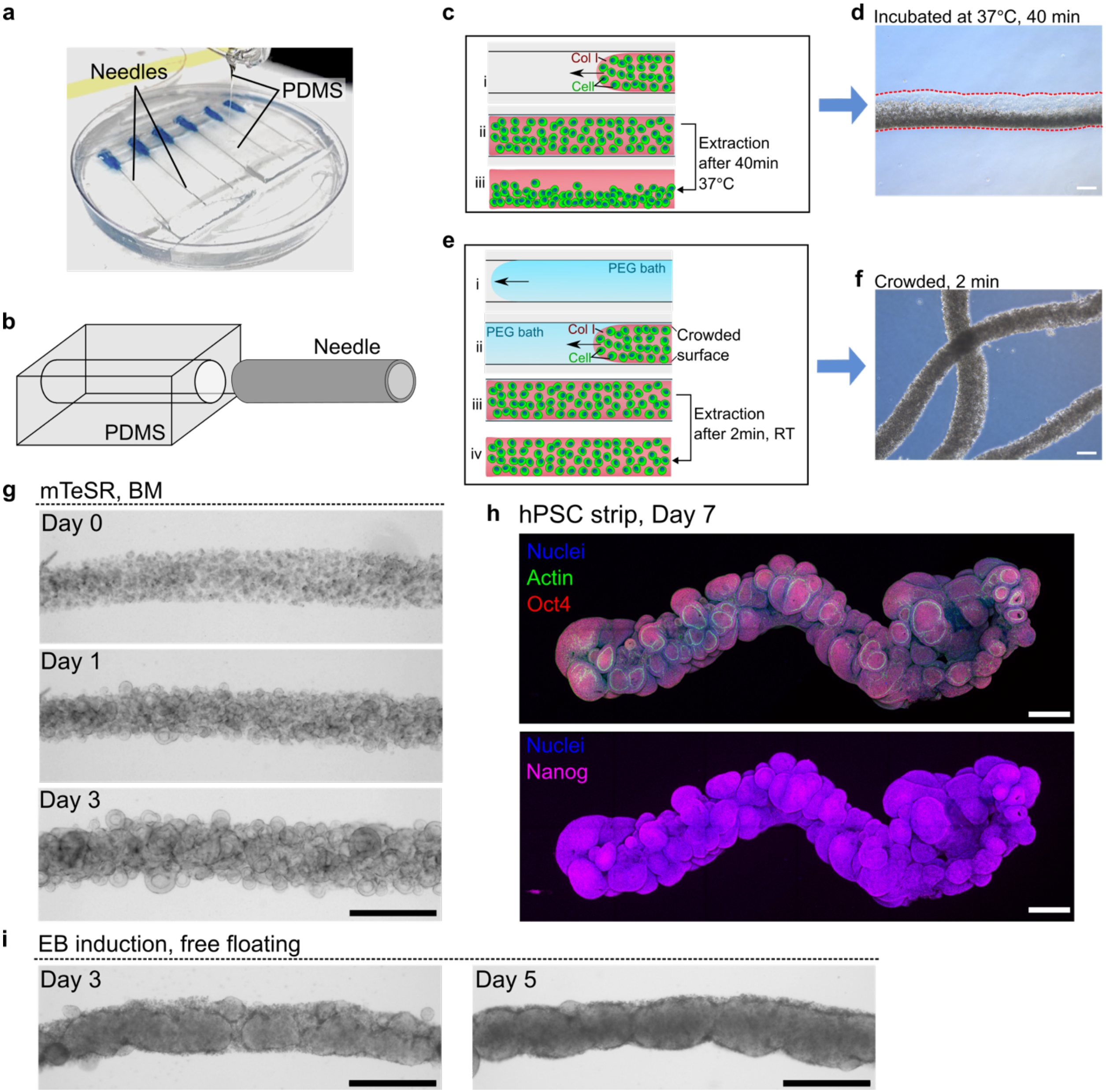


**Extended Data Fig. 5. Fabrication and culture of hPSC macroscopic scale strips.** (a) Macrograph and (b) schematic showing the fabricating procedure of the parallel channel device for rapid strip production. (c) Schematic of fabricating cell-laden collagen strip without PEG bath (i) high concentration of hPSCs ($4\times10^7$ cells/mL) incorporated neutralized collagen solution (1.5mg/mL) is injected through a PDMS molded channel. (ii) The strip is allowed to gel completely by incubating in the CO2 incubator at 37°C for 40 minutes and then flushed out by PBS shown in (iii). (d) Slow gelation of the collagen results in nonhomogeneous cell distribution within the stripe, indicating cells sedimented unevenly within the gel during the incubation process. (e) Schematic of fabricating cell-laden collagen strip with PEG bath. (i) A 200mg/mL PEG 8000 solution is first injected through the parallel channel, followed by the same bioink containing hPSC ($4\times10^7$ cells/mL) and 1.5mg/mL collagen solution shown in (ii). (iii) The collagen strip is allowed to stay in the channel at room temperature for only 2 minutes and then (iv) extracted. (f) Rapidly solidified cell-laden collagen strips exhibited reproducible homogeneous cell distribution. Scale bars in (d) and (f), 200 μm. (g) Bright-field images tracking the expansion of an hPSC strip within the Geltrex encapsulation. Over time, the strip exhibited pronounced outgrowth of organized, luminal colonies. Scale bar, 500 μm. (h) The hPSC strips were allowed to expand for up to 7 days. Immunofluorescent staining of F-actin, Oct4, and Nanog demonstrate hPSC well preserved their proper apical/basal polarity and pluripotency after the long-term culture. Scale bars: 500 μm. (i) Bright-field images showing EB induction of an hPSC strip freely cultured in EB induction medium on Day 3 and Day 5. The strip exhibited a solid structure and brightened surface cells. Scale bars, 500 μm.

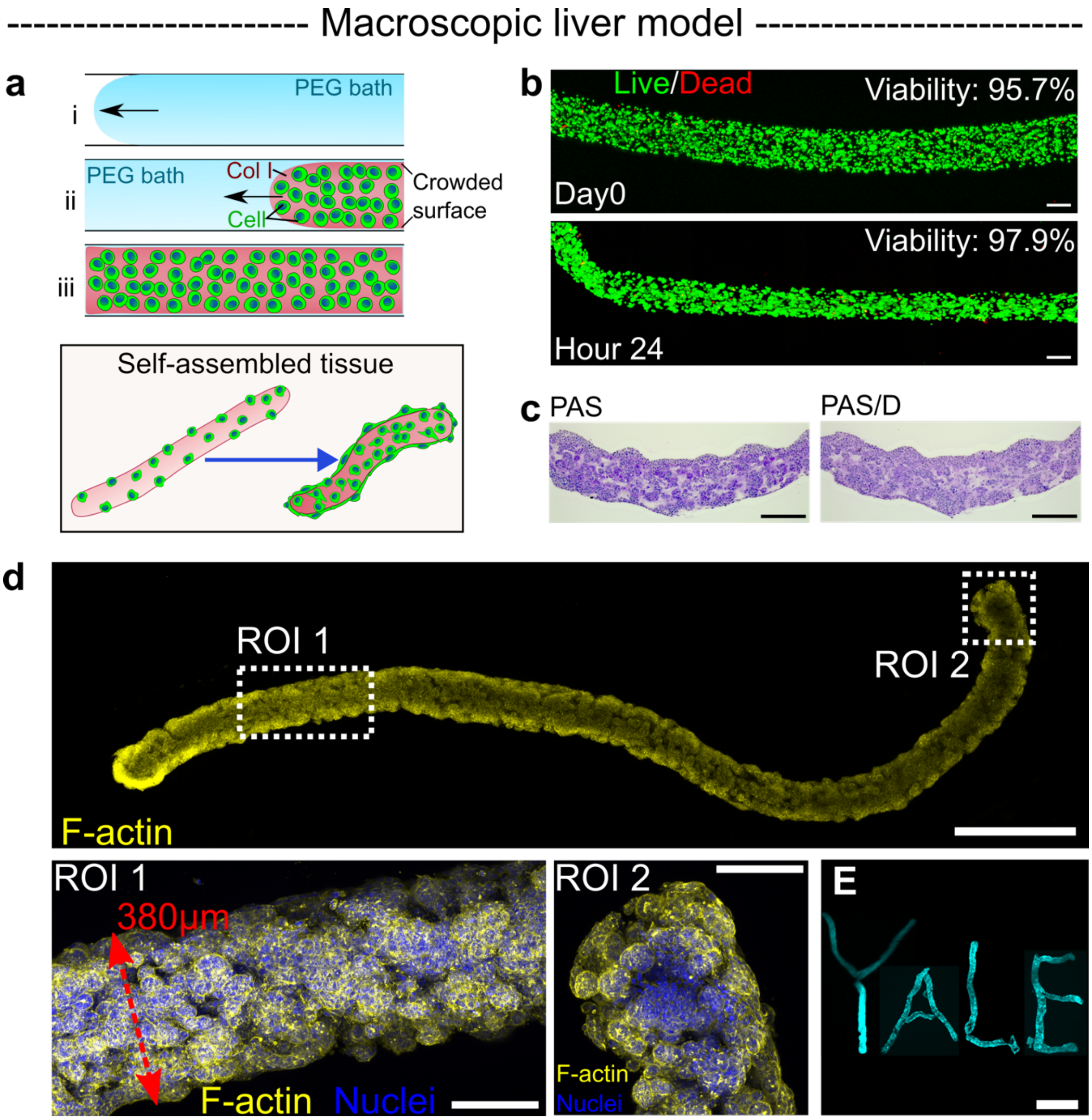


**Extended Data Fig. 6. Macroscopic liver cell constructs.** (a) Schematic of fabricating cell-lade macroscopic strips. (b) Viability assay on HepG2 strips after fabrication (Day 0) and after 24-hour culture. Scale bars, 200μm. (c) Histological staining of PAS and PAS with diastase (PAS/D) on a fixed 8-day HepG2 strip. Scale bars, 200 μm. (d) A HepG2 strip ($1\times10^7$ cell/mL) cultured freely in the medium for 8 days. Scale bar, 1mm. In the ROIs, actin and nucleus were stained. Scale bars, 200 μm. (e) Assembling the Day-7 HepG2 strips into a macroscopic complex pattern "YALE" in a culture medium-rich granular support. Scale bar, 2mm.

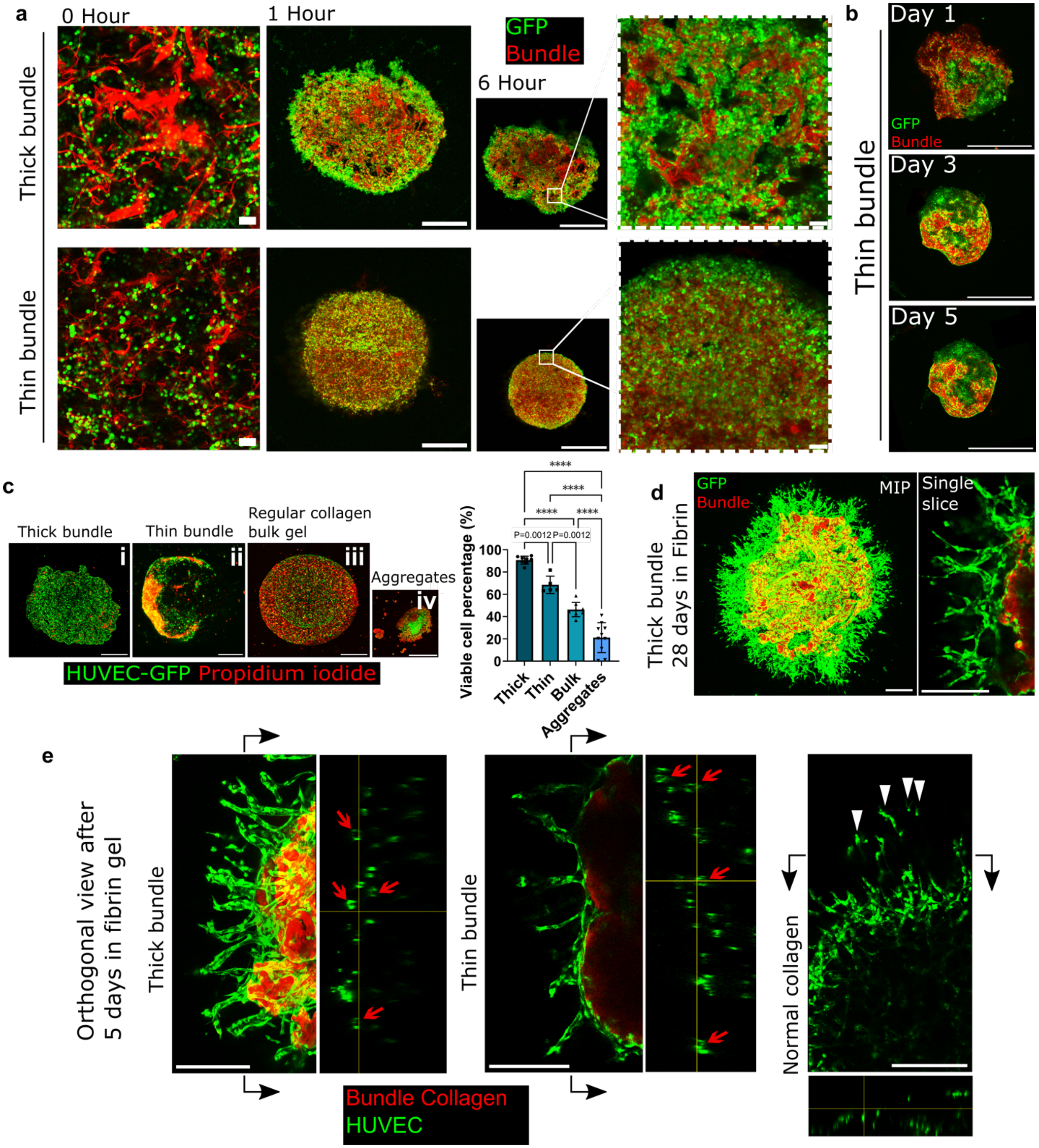


**Extended Data Fig. 7. Bundles guide EC patch formation and maintain long-term angiogenic sprouting.** (a) Live confocal images showing short tracking of EC patch formation within 6 hours. Images at Hour 0 showing loosely seeded HUVECs and μ-bundles. Scale bars, 100 μm. Images at Hour 1 and Hour 6 showing cell-μ-bundle self-assemble into patches within 6 hours. Scale bars, 1 mm. Scale bars in the magnified images at Hour 6, 100 μm. (b) Live confocal images showing daily tracking of patches with thin μ-bundles. Scale bars, 1000 μm. (c) Viability assays on HUVEC (i) growing on the thick μ-bundle scaffold (n=8; N=2); (ii) growing on thin μ-bundle scaffold (n=5; N=2); (iii) embedded in 2 mg/mL bulk collagen gels (n=8; N=2); and (iv) aggregated without any collagen (n=10; N=2), through dead cell staining propidium iodide (red). Scale bars, 500 μm. Bar plots represent mean ± S.D. ****$P < 0.0001$, **$P = 0.0012$, determined by one-way ANOVA, with Tukey's multiple comparisons. (d) Live confocal images showing long-term (28 days) angiogenic sprouting of thick μ-bundle patches. Scale bars, 300 μm. (e) Cross-sectional representations showing lumenal sprouts of EC-patches (red arrows) vs. single cell migration of 2 mg/mL regular bulk collagen gel (white arrowheads). Scale bars, 300 μm.

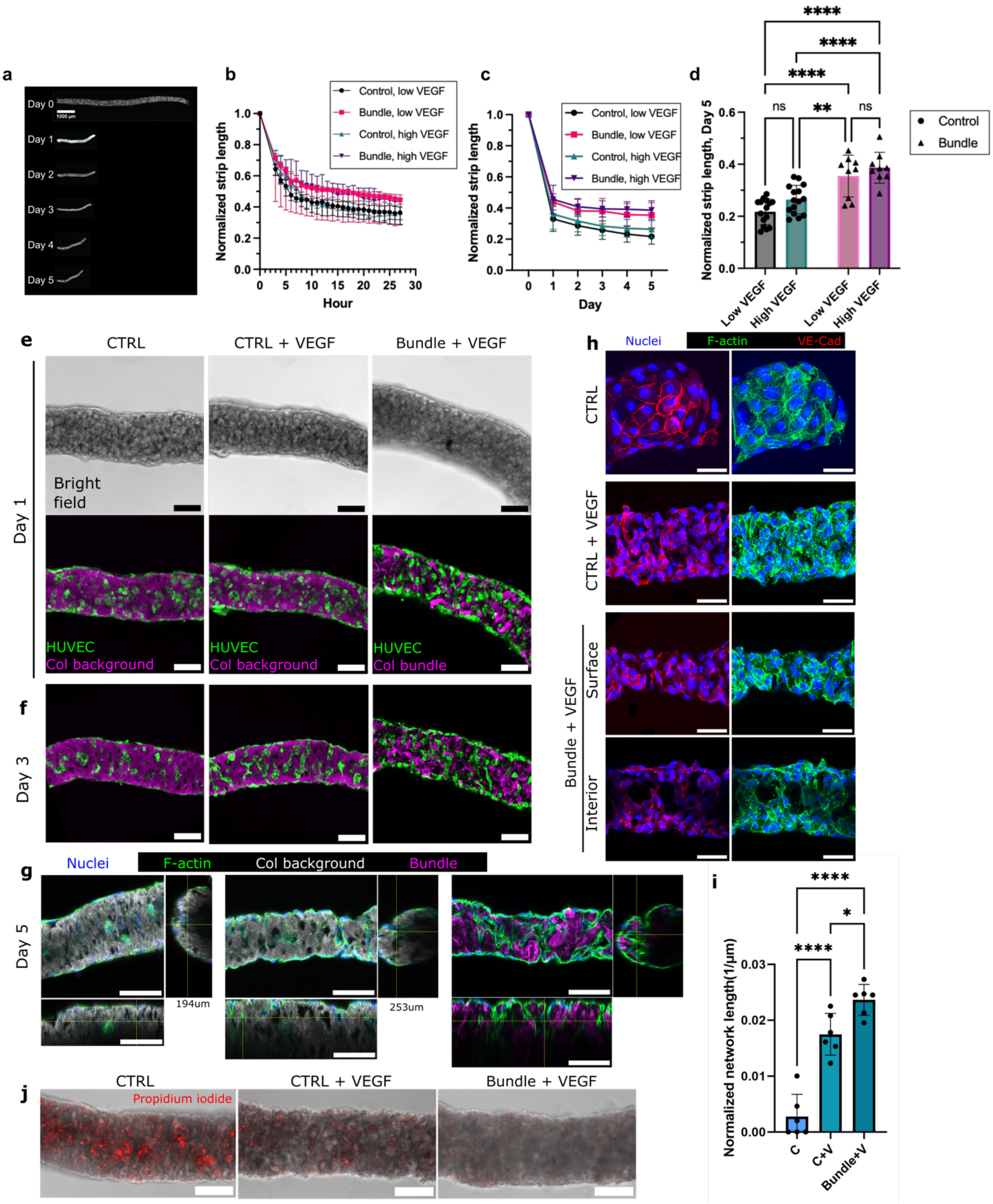


**Extended Data Fig. 8. Fabrication of EC macroscopic strips with μ-bundles.** (a) EC strip compaction. Scale bar, 1000 μm. Lengths of strips (mixed with or without bundles) normalized by Hour 0 were tracked within (b) hours and (c) days with low or high-dosing VEGF treatments. (d) Strip length comparison on Day 5. Two-way ANOVA with Tukey's multiple comparisons, ****P

< 0.0001, **P = 0.0043 (n=15 strips without bundles for both low and high VEGF conditions, n=9 strips with bundles for both low and high VEGF conditions, pooled from N=2-3 biological replicates). (e) Bright-field (upper row) and fluorescent confocal images (bottom row) of HUVEC strips, showing vascular constructs with strip geometry on Day 1. Scale bars, 100 μm. (f) HUVEC strip on Day 3. HUVECs undergoing guidance through μ-bundles. Scale bars, 100 μm. (g) Magnified confocal images and orthogonal views of strips on Day 5. μ-bundles incorporated with VEGF guide the interconnected network formation. HUVECs cover the strip surface, internal luminal structures are observed and connect to the surface HUVECs. Scale bars, 100 μm. (h) Magnified views of VE-Cad (red) and actin (green). Scale bars, 50 μm. (i) EC networks in strips are evaluated by normalized network length. (C, conditions without μ-bundles and without VEGF. C+V, conditions without bundles but with VEGF. Bundle+V, conditions with bundles and with VEGF.) Bar plots represent mean ± S.D. ****P < 0.0001, *P = 0.0221, determined by one-way ANOVA, with Tukey's multiple comparisons (n=14 or 13 from N=2 biological replicates). (j) Staining of dead cells with PI, demonstrating VEGF treatment and mesoporous architecture promote cell survival.

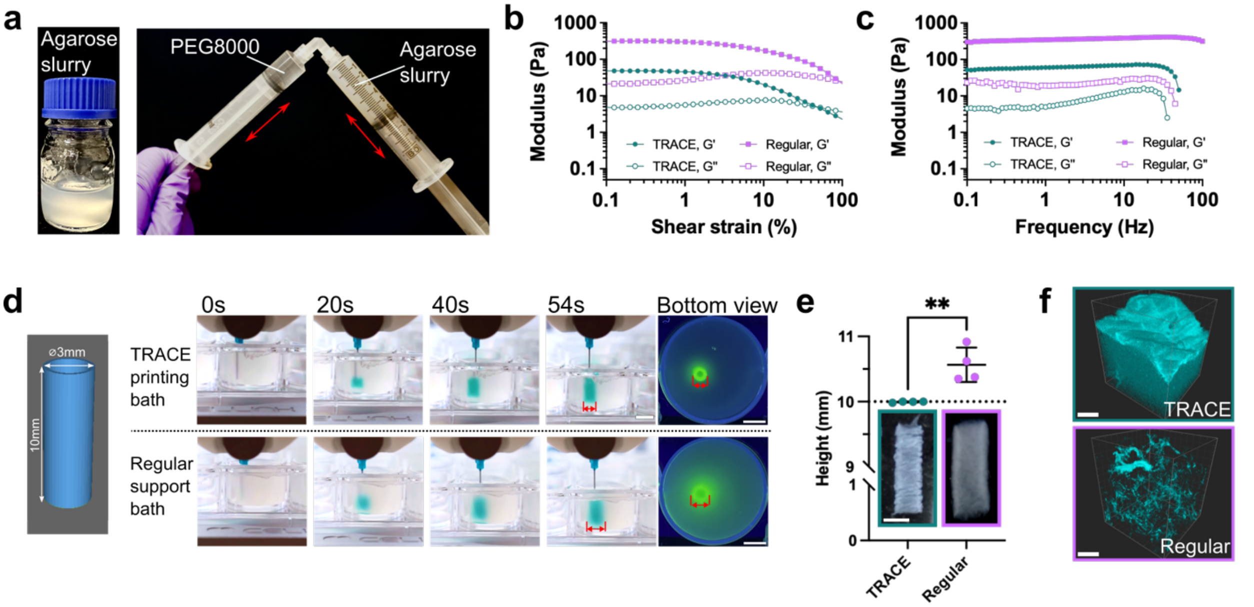


**Extended Data Fig. 9. Developing and printing in the TRACE bath.** (a) The procedure of preparing the TRACE printing bath by mixing agarose slurry (left) and PEG solution in two syringes connected with a 90-degree angled luer lock connector (right). In this study, we term the agarose slurry the "regular bath." (b) Rheological characterization of shear moduli (G': storage modulus; G'': loss modulus) of TRACE bath and regular bath as a function of shear strain at the oscillation frequency of 1Hz. (c) Shear moduli of TRACE bath and regular bath as the function of shear frequency at the strain of 0.1%. (d) Real-time monitoring of printing a tubular construct (diameter: 3mm; height: 10mm) vertically with a low viscosity collagen ink (2mg/mL, neutralized). The ink was loaded with blue food color and fluorescent microbeads to visualize the printing procedure and post-printing fidelity in the baths after a 30-minute incubation at room temperature. Note that the printed tubes seem broader in the side profile view due to the optical distortion from the curved wells. In the regular slurry, significant diffusion of ink and broadened diameter were observed. Scale bar in the printing process, 1cm; scale bars in the bottom view of the printed tubes, 5mm. (e) Height measurement of the released collagen tubes, showing tubes printed in the regular bath were elongated due to ink diffusion in the vertical direction (N=4 for each condition). A two-tailed student's t-test was performed (**P=0.005). Scale bar, 3mm. (f) 3D reconstruction of the surfaces of the collagen tubes printed in the TRACE bath vs. the regular bath, comparing the surface texture and porosity. Scale bar, 30 µm.

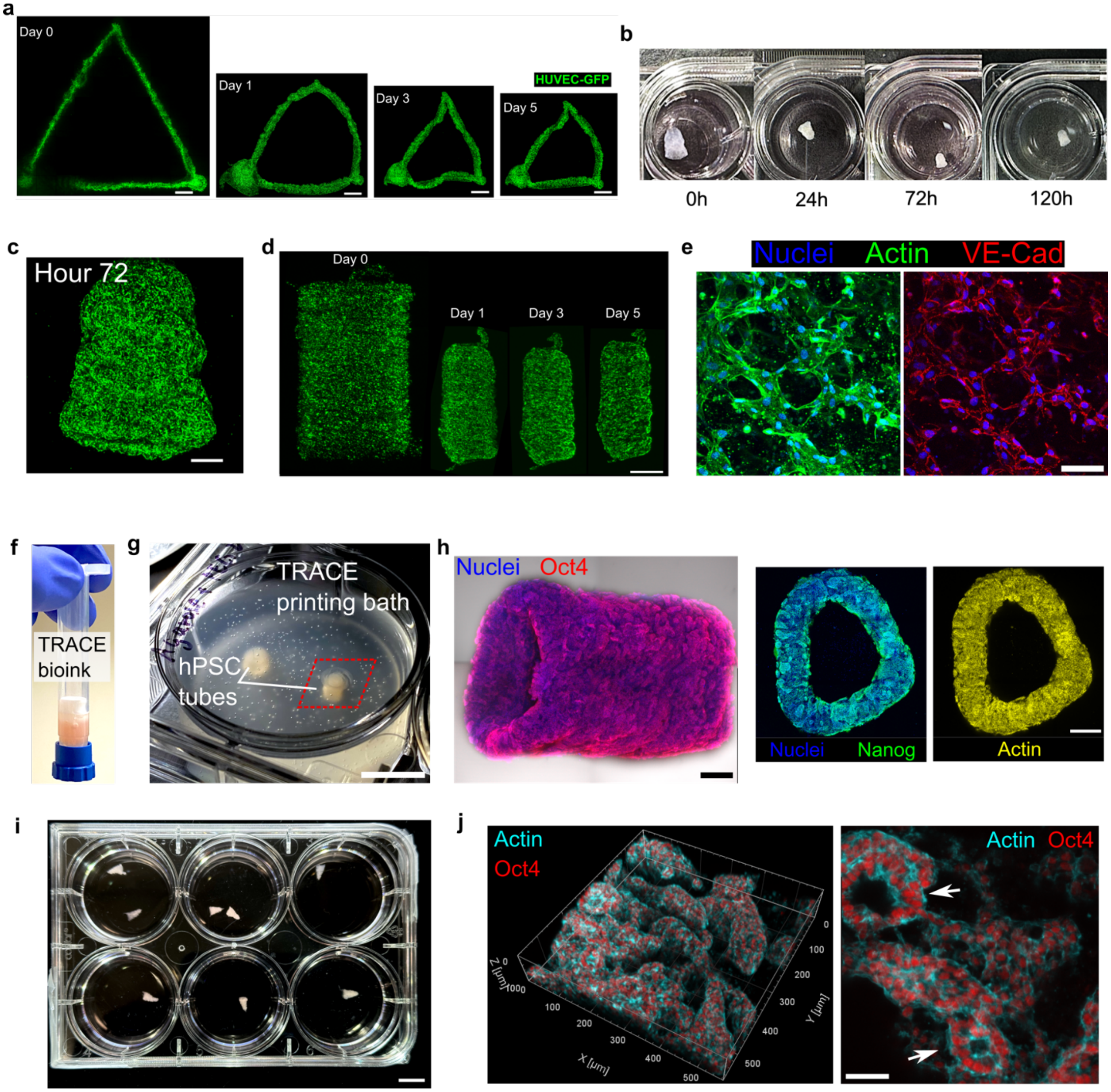


**Extended Data Fig. 10. Printing 2D and 3D bioactive multicellular tissues.** (a) TRACE printing of millimeter-scale triangles consisting of GFP-labeled HUVEC with low-concentration collagen (2mg/mL). Daily live confocal imaging showing the morphological evolution of the whole HUVEC triangular construct for 5 days. Scale bar, 2mm. (b) TRACE printing of HUVEC-collagen bioink into a 3D bell-shaped construct cultured in the medium for 120 hours (5 days) in a 24-well low attachment plate. Fluorescent imaging of the whole tissue (GFP) on Day 3 (Hour 72) is shown in (c). Scale bar, 500 µm. (d) TRACE printed HUVEC tube ($4\times10^6$ cells/mL) with neutralized 2mg/mL collagen ink, and was allowed to grow for 5 days. Scale bar, 1mm. (e) Fluorescent staining of actin and VE-Cad showing the HUVEC organization on the printed surface. Scale bar, 100 µm. (f) High cell concentration hPSC bioink in low concentration collagen (2mg/mL) was used to print (f)-(h) tubular structures in the TRACE support bath. (g) Macrograph showing hPSC tubes printed in the TRACE bath. 10mm. (H)Confocal imaging shows the side view and the cross-sectional view of the hPSC tube with staining of pluripotency markers and actin. Scale bars, 500μm. (i) High-throughput production of printed hPSC ventricles demonstrates the reproducibility of TRACE printing with cell-based collagen ink. Scale bars: 10mm. (j) Close-up imaging on the surface of the hPSC tube showing the luminal cellular organization with F-actin staining (indicated by white arrows) and pluripotency labeled by Oct4. Scale bar, 50μm.

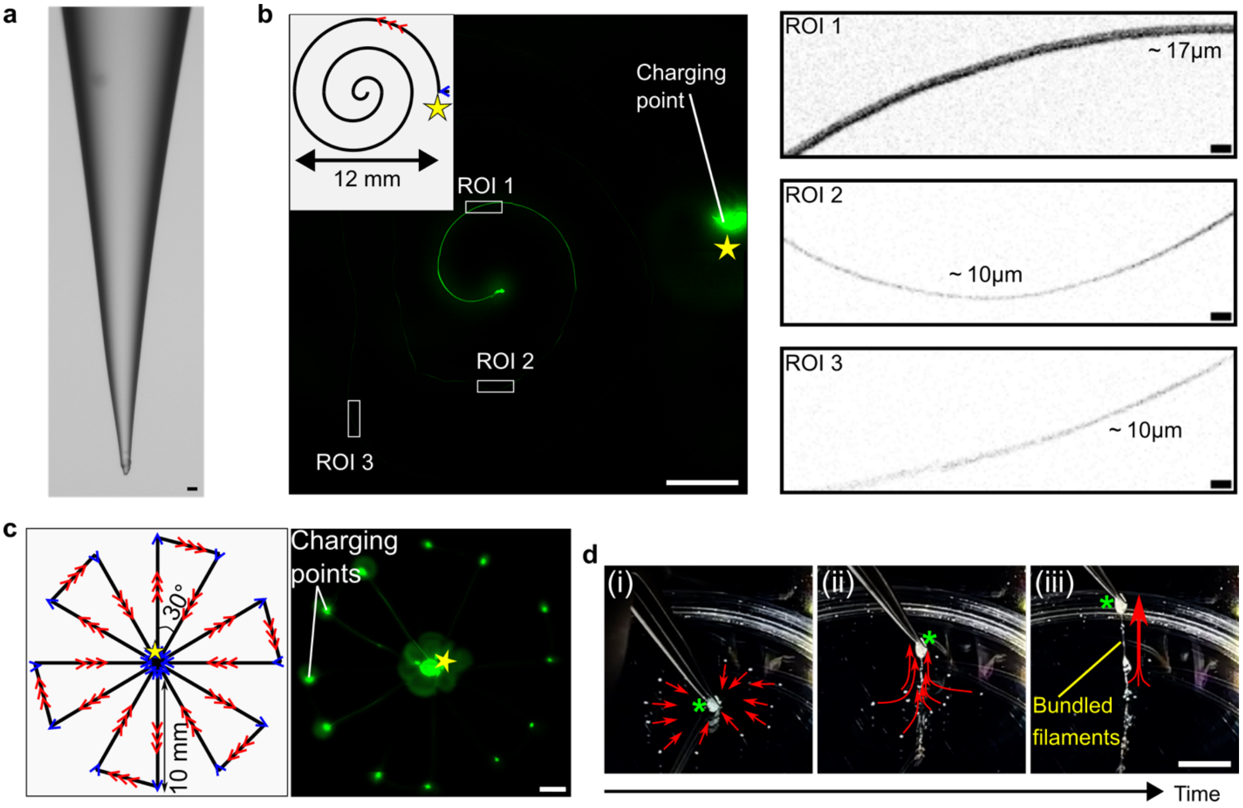


**Extended Data Fig. 11. Complex patterning of collagen ultrathin filaments.** (a) Pulled glass nozzle for ultrathin collagen filament patterning. Scale bar, 20 µm. (b) Patterning of ultrathin collagen filament (10-17 µm) into a spiral shape. Scale bar in the global pattern, 2mm. Scale bars in the ROIs, 50µm. (c) A windmill pattern of ultrathin filaments (printing speed: 4,000mm/min) connecting multiple material charging points as more visible landmarks for the printed pattern. Scale bar, 2mm. For (b) and (c), the stars indicate the starting point of the printing paths. The directions of the arrows indicate the paths' direction. The red multiple arrows indicate high-speed paths, and the single blue arrows indicate low-speed paths. (d) Structural folding of the windmill pattern (Movie. 10) by pulling up the center (indicated by the green asteroid). Upon pulling, the movements of the charging points indicate these more visible components are tethered to the center by the ultrathin filaments. This mechanism can be used to collect and bundle up multiple printed ultrathin filaments. Scale bar, 5mm.

| **Collagen pre-bundle concentration** | **2 mg/mL*** | **4 mg/mL** | **6 mg/mL** | **8 mg/mL** | **9.5 mg/mL*** |
|---|---|---|---|---|---|
| Cell water (μL) | 140.2 | 100.5 | 61.2 | 21.5 | 0 |
| 10× PBS (μL) | 20 | 20 | 20 | 20 | 0 |
| 9.5 mg/mL acidic collagen (μL) | 38.1 | 76.2 | 114.3 | 152.4 | 200 |
| NaOH (μL) | 1.7 | 3.3 | 4.5 | 6.1 | 0 |
| Total volume of collagen precursor (μL) | 200 | 200 | 200 | 200 | 200 |
| Bundle re-suspension volume (μL) | 40 | 80 | 120 | 160 | 200 |
| **Final collagen concentration of bundle re-suspension (mg/mL)** | **9.5** | **9.5** | **9.5** | **9.5** | **9.5** |

*For the self-assembled EC-patches, we term bundles made with 2mg/mL collagen precursor as "thin" bundles, and term bundles made with 9.5mg/mL collagen precursor as "thick" bundles.

**Extended Table 1. Recipe for bundle re-suspension made from collagen precursors with varying concentrations.**

| Differentiation Timing | Reagents |
|---|---|
| Day 0<br>(cell confluence >98%) | mTeSR and RPMI/B27 minus insulin (volume ratio of mTeSR and RPMI/B27 minus insulin is 1:3; RPMI/B27 minus insulin contains RPMI and B27 minus insulin at a ratio of 49:1), containing 12.5 μM CHIR99021 |
| Day 1<br>(after 24 hours) | RPMI/B27 without insulin |
| Day 3<br>(Same hour of the day as adding CHIR99021) | RPMI/B27 without insulin, containing 5 μM IWP4 |
| Day 5<br>(Additional 48 hours) | RPMI/B27 without insulin |
| Day 7 | RPMI/B27 without insulin |
| Day 9 | RPMI/B27 |
| Day 11 | RPMI/B27 |
| Day 12 | RPMI without glucose, containing 4mM lactate |
| Day 14 | RPMI without glucose, containing 4mM lactate |
| Day 16 | RPMI/B27 |
| Day 18 | Ready to seed |

**Extended Table 2. Cardiac differentiation from hPSCs**

**Movies**

**Movie 1.** High throughput, rapid formation of micro-liter collagen disks. Neutralized, 2mg/mL collagen droplets (5 μL) are dispensed onto the PEG8000 bath (200mg/mL in PBS) and form defined disk shapes in seconds with increasing opacity.

**Movie 2.** Micro-liter collagen droplets dispensed into PBS. Neutralized, 2mg/mL collagen droplets (5 μL) are dispensed onto the PBS and instantly disappear. No visible gelation is observed within 5 minutes. Around Minute 6, some droplets start to form gels with undefined shapes.

**Movie 3.** Rapid fabrication of collagen strip in the channel device. Neutralized collagen solution is injected from the left side into a PDMS channel prefilled with PEG8000 solution. Distinct interface between collagen solution and PEG bath can be observed.

**Movie 4.** Rapid single line printing of collagen ink (neutralized, 2mg/mL) in the TRACE printing bath vs. the regular agarose granular bath. The TRACE bath enables rapid gelation of the deposited collagen ink and fast extraction (1 minute), whereas the printed collagen does not solidify in the regular granular support bath and cannot be extracted from the bath 1 or 5 minutes after printing.

**Movie 5.** Rapid releasing of 3D collagen construct printed in the TRACE bath. A millimeter-scale 3D hollow gourd shape printed with acidified collagen ink (6mg/mL) in the TRACE bath shows expedited gelation at the room temperature and can be released from the bath in 3 minutes. Room temperature for 3 minutes in the regular support bath (without PEG) is not sufficient to lead to full gelation and structural integrity upon releasing.

**Movie 6.** Rapid releasing of a 3D collagen tube (diameter: 3mm, height: 5mm) printed in the TRACE bath. The millimeter-scale collagen tube printed with acidified collagen ink (6mg/mL) in the TRACE bath gels at the room temperature and can be released from the bath in 3 minutes.

**Movie 7.** High-speed (4000mm/min) printing of a macroscopic windmill pattern (2cm in diameter) consisting of microscopic ultrathin collagen filaments and material charging points in PEG8000 bath.

**Movie 8.** Structural folding of a macroscopic windmill pattern (diameter: 2 cm) consisting of ultrathin collagen filament. The visible charging points are connected to the central point by the ultra-thin filaments and can be bundled and maneuvered by pulling only the central point.

**Movie 9.** Whole-tissue spontaneous calcium and beating of a TRACE-printed cardiac ventricle construct. The calcium signal and tissue contraction are synchronized.

**Movie 10.** Cardiac cycles of a TRACE-printed cardiac ventricle construct. (Left: PIV of the fluid field; Right: recording of the fluid output visualized by fluorescent beads.)
